# A Non-stop identity complex (NIC) supervises enterocyte identity and protects from pre-mature aging

**DOI:** 10.1101/2020.08.23.263095

**Authors:** Neta Erez, Lena Israitel, Eliya Bitman-Lotan, Wing Hing Wong, Gal Raz, Salwa Danial, Na’ama Flint Brodsly, Elena Belova, Oksana Maksimenko, Pavel Georgiev, Todd Druley, Ryan Mohan, Amir Orian

## Abstract

A hallmark of aging is the inability of differentiated cells to maintain their identity. In the aged *Drosophila* midgut differentiated enterocytes (ECs) lose their identity, and the integrity of the midgut tissue and its homeostasis are impaired. To discover regulators of EC identity relevant to aging we performed an RNAi screen targeting 453 ubiquitin-related genes in fully differentiated ECs. Seventeen genes were identified, including the de-ubiquitinase Non-stop (Not/dUSP22; CG4166). Acute loss of Non-stop in young ECs phenotypically resembled aged ECs. Lineage tracing experiments established that Non-stop-deficient young ECs as well as wild-type aged ECs are no longer differentiated. Aging or acute loss of Non-stop also resulted in progenitor cell hyperproliferation and mis-differentiation, loss of gut integrity, and reduced organismal survival. Proteomic analysis unveiled that Non-stop maintains identity as part of a Non-stop identity complex (NIC) that contains E(y)2, Sgf11, Cp190, (Mod) mdg4, and Nup98. Transcriptionally, Non-stop ensured chromatin accessibility at EC genes, maintained an EC-specific gene expression signature, and silenced non-EC-relevant transcriptional programs. Within the NIC, Non-stop was required for stabilizing of NIC subunits. Upon aging, the levels of Non-stop and NIC subunits declined, and the large-scale organization of the nucleus was distorted. Maintaining youthful levels of Non-stop in wildtype aged ECs safeguarded the protein level of NIC subunits, restored the large-scale organization of the differentiated nucleus, and suppressed aging phenotypes and tissue integrity. Thus, the isopeptidase Non-stop, and NIC, supervise EC identity and protects from premature aging.

## Introduction

Differentiated cell states are actively established and maintained through action of “identity supervisors” (Holmberg & Perlmann, 2012; Natolli, 2014). Identity supervisors control expression of genes that enable differentiated cells to respond to environmental cues and perform required physiological tasks. Concomitantly, they ensure silencing/repression of previous fate and non-relevant gene programs and reduce transcriptional noise. Inability to safeguard differentiated cell identity is a hallmark of aging and results in diseases such as neurodegeneration, diabetes, and cancer (Bensellam et al. 2018; Hudish, et al. 2019; Conway et al 2015; Hnisz et al. 2013; Deneris & Hobert 2014). In many cases, transcription factors (TFs) together with chromatin regulators and architectural/scaffold proteins establish and maintain large-scale chromatin and nuclear organization that is unique to the differentiated state of the cell (Blau & Baltimore 1991; Booth & Brune 2016; Naetar, et al 2017; Bitman-Lotan & Orian 2018).

In adult *Drosophila* midgut epithelia, the transcription factor Hey (Hairy/E(spl)-related with YRPW motif), together with *Drosophila* nuclear type A lamin, Lamin C (LamC), co-supervise identity of fully differentiated enterocytes (ECs) (Monastirioti et al. 2010; Gruenbaum and Foisner R. (2015) Flint-Brodsly et al. 2019). Highly similar to vertebrate gut, *Drosophila* midgut epithelia intestinal stem cells (ISC) either self-renew or differentiate into progenitor cells that mature into enteroendocrine cells (EEs) or give rise to enteroblast progenitors. Enteroblasts (EB) mature into fully polyploid differentiated enterocytes (ECs) that carry out many critical physiological tasks of the intestine (Figure 1A; Jiang, & Edgar 2012; Lemaitre & Miguel-Aliaga 2013; Buchon et al. 2013; Hung et al. 2020). Aging affects the entire midgut, and is associated with loss of EC identity, mis-differentiation of progenitors, pathological activation of the immune system, and loss of the physiological properties of the gut and its integrity. It also results in loss of intestinal compartmentalization, and microbiota-dysbiosis, all leading to reduced lifespan (Biteau et al. 2010; Rera et al 2012; Bonnay et al. 2013; Ferrandon, 2013; Chen et al. 2014; Li, et al. 2020; Rodriguez-Fernandezet al 2020; Jasper H. 2020). During aging, the protein levels of identity supervisors such as Hey and LamC decline, resulting in inability to maintain EC-gene programs and ectopic expression of previous- and non-relevant gene programs (Neves et al 2015; Takeda et al 2018; Flint-Brodsly et al. 2019). Indeed, continuous expression of Hey in aged ECs restores and protects EC identity, gut integrity, and tissue homeostasis (Flint-Brodsly et al. 2019).

**Figure 1:**
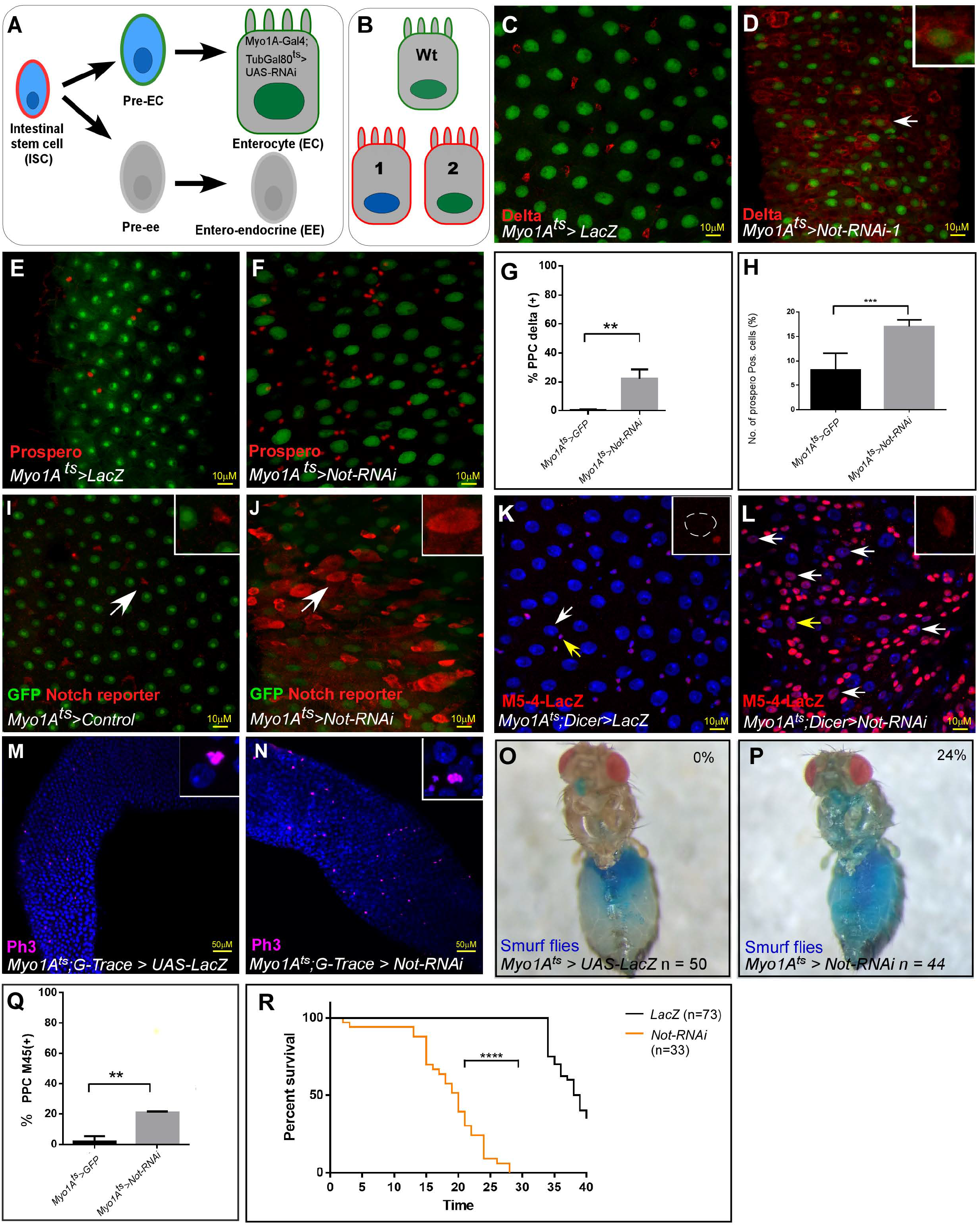
RNAi Screen identified Non-stop (Not) as an ECs identity supervisor. **(A)** Schematic diagram of midgut differentiation and an outline of the Ub/UbL screen (see text for details). The Notch ligand, Delta, is expressed on the surface of Intestinal stem cells (ISC) marked in red. **(B)** Phenotypes expected from positive hits: 1. Loss of expression of EC-specific GFP (expressed only in fully differentiated ECs using MyoIA>Gal4/Gal80^ts^ system), along with ectopic expression of the ISC marker Delta (red). 2. Polyploid cells that ectopically express Delta and retain expression of GFP. **(C-F)** Confocal images using UAS-LacZ (C, E) or UAS-Not RNAi (D, F) along with UAS-GFP expressed under the control of MyoIA>Gal4/Gal80^ts^ system. Scale bar is 10μM. The stem cell marker Delta (C, D) and EE marker Prospero (E, F) are shown in red. **(G, H)** Quantification of three biological repeats of experiments similar to that shown in C-F. \***(I, J)** Expression of UAS-Not RNAi, but not control, in ECs for 48h using MyoIA>Gal4/Gal80^ts^ results in ectopic expression of the Notch-reporter (red) in polyploid cells. **(K, L, Q)** Expression of the *escargot* progenitor enhancer reporter M5-4-LacZ in control or Not-targeted ECs (red). Yellow arrows points to cells shown in the insets. White arrows in L are examples of EC-like polyploid cells ectopically expressing the reporter. **(M, N)** Loss of Not in ECs resulted in an increase in the mitosis marker p-H3 in small cells. **(O, P)** Loss of Not in ECs impairs gut integrity as evident by the leakage of blue-colored food into the abdomen (smurf assay); 24% of Not-RNAi flies show loss of gut integrity versus 0% in control flies (n = 44, 55 respectively, p<0.001) **(Q)** Quantification of M5-4 positive PPCs in control and upon targetnig Not in ECs. (***= p<0.001 **p<0.01). **(R)** Survival analysis of flies expressing the indicated transgenes in ECs under the control of MyoIA-Gal4/Gal80ts (****=p<0.0001).

Regulation of EC identity requires signaling to the nucleus to communicate physiological changes in the gut environment. An important dynamic signaling mechanism involves changes in post-transcriptional modifications which may propagate, amplify, or conduct signals, ultimately leading to differential gene regulation. One type of posttranslational modification is the covalent attachment of ubiquitin or ubiquitin-like (Ub/UbL) molecules, that affect protein stability, function, localization, as well as modulate chromatin structure (Heideker, Wertz; 2015; Swatek & Komander, (2016); Cappadocia, & Lima, 2018; Song & Luo, 2019; Yao et al. 2020). Recent works suggest an intimate links between ubiquitin proteostasis and aging (Kevei, & Hoppe, 2014; Vilchez et al. 2014; Hohfled & Hoppe, 2018; Enam et al 2018; Chua & Signer, 2020). Therefore, we performed an RNAi screen to search for Ub/UbL-related genes within ECs that supervise identity. Screening 548 genes, 17 were identified whose conditional elimination in fully differentiated ECs resulted in loss of EC identity. Further analysis revealed one of them, the deubiquitinating isopeptidase (DUB) Non-stop (Non-stop/dUSP22) is a key EC identity supervisor. Purification and proteomic analysis identified Non-stop as part of a CP190/Nup98/Sgf11/e(y)2/mdg4 protein complex, termed Non-stop identity complex (NIC), that is essential for maintenance of EC identity. In part, Non-stop protects NIC proteins from age-dependent decline, safeguarding the EC-gene expression signature, as well as large-scale nuclear organization in these cells, preventing premature aging. Over lifespan, Non-stop protein levels in ECs declined, leading to loss of NIC subunits. This decline is associated with loss of gut identity and physiology at the cellular and tissue levels and maintaining youthful levels of Non-stop prevented loss of the NIC and prevented aging of the gut.

## Results

### A transgenic RNAi screen identified Ub/UbL-related EC identity regulators

To identify EC identity supervisors, a collection of RNAi transgenic flies targeting 453 evolutionarily conserved Ub/UbL-related genes were screened (Table S1; List of Ubiquitin-Related Genes according to DRSC - http://www.flyrnai.org/DRSC-SUB.html). Genes were knocked-down in fully differentiated ECs of 2-4 day old adult *Drosophila* females using UAS-RNAi lines and EC-specific conditional driver *MyoIA*-Gal4/Gal80^ts^ coupled system (termed MyoIA^ts^; see methods for specific lines used; Salmeron et al., 1990; Brand and Perrimon, 1993; Flint Brodsly et al. 2019). Conditional RNAi was achieved by shifting flies from 25 to 29^0^C for 48 hours, after which guts were dissected and analyzed. Immunofluorescence was used to score loss of proteins which are hallmarks of EC identity (Figure 1A, B). Among the changes upon loss of EC identity is the as ectopic expression of the ISC marker Delta on the surface of ECs-like polyploid cells. This change may be also accompanied with a decline in the expression of a GFP signal expressed only in fully differentiated ECs (derived from the MyoIA^ts^ Gal4, UAS-GFP transgene). Knockdown of seventeen genes resulted in loss of EC identity and the appearance of EC-like polyploid cells (PPCs). We also determined whether these genes are required for maintaining identity of enteroendocrine cells (EE’s), or progenitor cells, using the Prospero>Gal4/Gal80^ts^, or Esg>Gal4/Gal80^ts^ that activates the UAS-RNAi in these cells, respectively (Figure 1, Supplemental Table S1).

### Non-stop supervises EC identity

Among the genes identified were E3 ubiquitin ligases, E2 enzymes, SUMO-related enzymes and ubiquitin-specific peptidases (DUB/USPs). We also identified few nuclear proteins harboring PHD domain that serve as binding to methylated histones, but may confer ubiquitin ligase activity present in established ubiquitin ligases (Examples are shown in Figure1, Supplemental Figure 1A-H,). The entire results of the screen are detailed in Figure 1, Supplemental Table S1 section of the table and validated positive hits under secondary screen). The DUB Non-stop (Not, dUSP22, CG4166) was identified as a bona-fide EC identity supervisor. RNAi mediated knockdown of Non-stop in ECs using three independent UAS-RNAi lines resulted in the inability of ECs to maintain MyoIA>UAS-GFP signal followed by ectopic expression of the ISC marker Delta on the surface of EC-like polyploid cells in both females and males (Figures 1C, 1D, quantitated in 1G and Figure 1 Source-data; Figure 1 Figure supplemental 1I-K).

Originally, Non-stop was discovered as a ubiquitin protease essential for axonal guidance in the visual system (Martin et al 1995). Non-stop is highly conserved from yeast to humans (Ubp8 and USP22 respectively; Mohan et al. 2014b), and its activity is required for deubiquitinating monoubiquitinated histone H2B (H2Bub) and activating gene expression (Weake et al. 2008; Mohan et al. 2014; Morgan et al 2016).

Immunofluorescence revealed that Non-stop is expressed in all gut cells (Figure 1 Supplemental Figure 2A-D). Non-stop is the major H2Bub deubiquitinase in *Drosophila,* therefore functional loss of Non-stop should lead to a many-fold increase in H2Bub levels (Weake et al. 2008; Zhang et al 2008; Morgan et al. 2016; Mohan et al, 2014; Li et al. 2017). Analysis showed that ECs lacking Non-stop exhibited more H2Bub, and accordingly protein extracts derived from these midguts were characterized by an over 6-fold increase in H2Bub compared to control knockdown midguts (Figure 1 Supp. Figure 2F-H). However, reduction of Non-stop in EEs did not impact EE or EC identity or number, or Delta expression, indicating that Non-stop function in maintaining gut identity was specifically localized in ECs (Figure 1 Supp. Figure 2I-L).

Non-stop knockdown in ECs resulted in an increase in Prospero positive cells (likely EEs) (Figure 1E, F, and quantified in 1H, Figure 1 Source-data). Non-stop elimination also affected the entire midgut tissue; resulting in ectopic activation of the Notch pathway, as well as the stem-cell enhancer M5-4 *esg::LacZ* in polyploid cells, indicating that these cells were losing differentiated state (Figure 1I-L, 1Q, and Figure 1source data). Loss of Non-stop also resulted in increased phospho-Histone H3 which is indicative of mitotic activity in small cells, likely progenitors (Figure 1M, N).

At the tissue level, knockdown of Non-stop in ECs reduced epithelial integrity as evidenced by leaking of blue colored food outside the gut, and reduced overall survival (Figure 1O, P and 1R respectively; Figure 1 source data).

We evaluated the identity and fate of young ECs conditionally lacking Non-stop, as well as aged wildtype ECs, as well as the cellular composition of the gut under these conditions. Towards this end we used the lineage tracing system G-TRACE. G-TRACE is a dual-color GAL4-dependent system, that enables tracing fully differentiated non-dividing cells (Figure 2A; Evans et al. 2009; Flint-Brodsly et al. 2019). In brief, the MyoIA-Gal4/Gal80^ts^ directs the expression of the color system only to fully differentiated ECs. Expression of the Gal4 is dependent on the EC-specific MyoIA promoter that induces expression of a UAS-RFP (red). Concomitantly, this Gal4 activity induces the expression of a Flp-recombinase resulting in a recombination event that drives permanent expression of a GFP regardless of the differentiation state of the cell. Therefore, wildtype young ECs express both RFP and GFP, and are the only population of polyploid cells (PPCs), observed in control midgut tissue (Figure 2B). In contrast, in guts where Non-stop was knocked-down in ECs, other populations of fluorescently colored PPC’s were observed. These include PPCs that express only GFP, termed PPC** (PPC^GFP+RFP-^; Figure 2C, and quantitated in 2H; Figure 2 source data). Unlike control cells, PPC** did not express EC-related transcription factors such Odd-skipped (Figure 2D-E). They also exhibited reduced expression of the differentiated lamin, LamC, (Figure 2F, G). By the nature of the G-TRACE system, we concluded that these PPC** are likely ECs that were no longer fully differentiated, failing to maintain EC identity. In accordance, PPCs that did not express EC key transcription factors such as Pdm1 and caudal were also observed (Figure 2 Figure Supplemental 1A-D). In addition, guts where Non-stop was targeted in ECs were populated with PPCs lacking expression of either RFP or GFP (termed PPC*), and are likely misdifferentiated progenitors that failed to activate the *myo-promoter* and the entire RFP/GFP marking system (Figure 2H, Figure 2 Source data).

**Figure 2:**
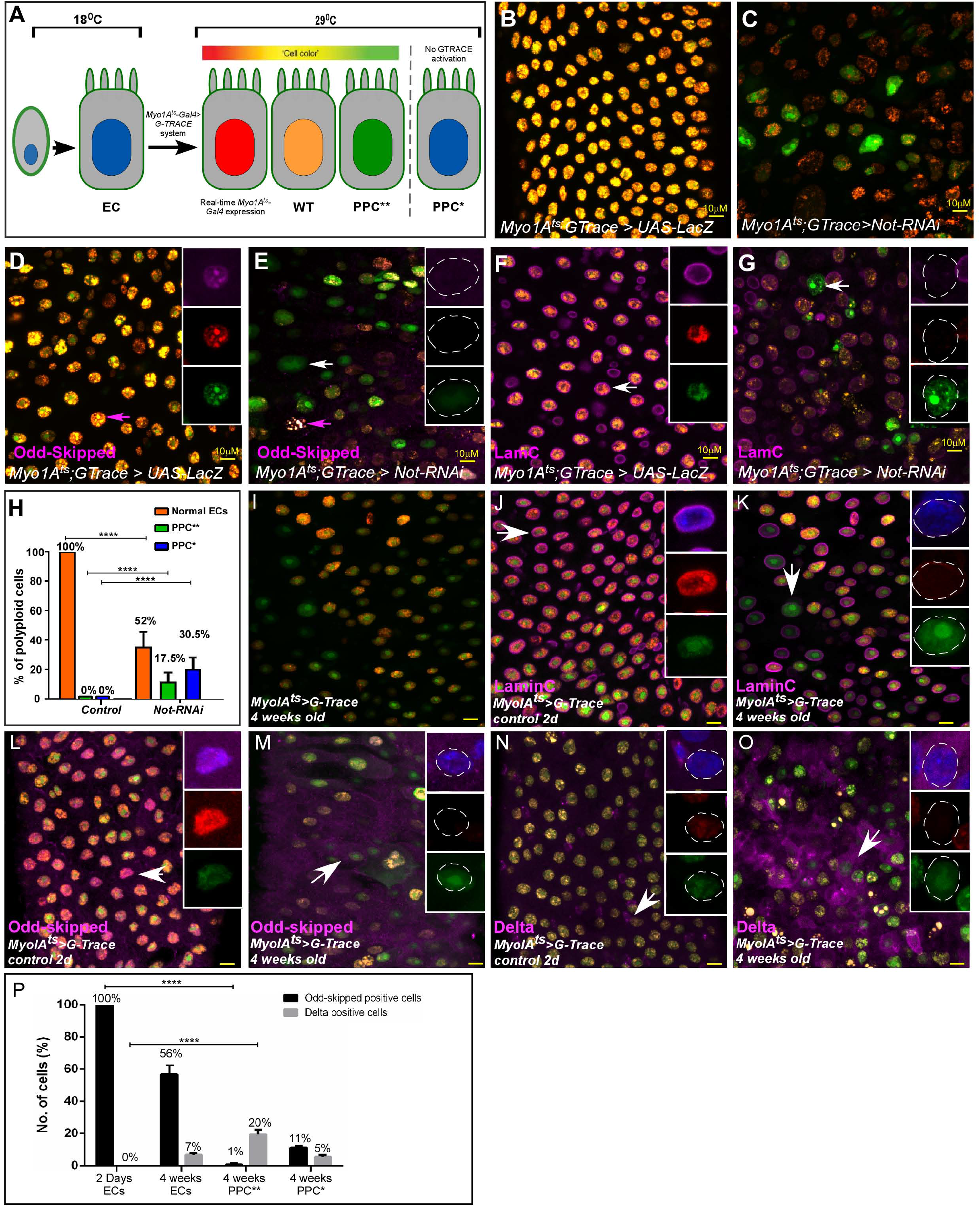
G-TRACE-Lineage characterization of Not targeted young ECs and, aged ECs. **(A)** Schematic diagram of EC-G-TRACE-lineage tracing system adopted from (Flint-Brodsly 2019). PPC** (RFP-GFP^+^), are EC that are no-longer differentiated. PPC* (RFP-GFP-) are miss-differentiated progenitors. **(B-O)** Confocal microscopy of midguts expressing the indicated transgenes, under the control of MyoIA^ts^ G-TRACE system using the indicated antibodies. DAPI (blue) marks DNA. Arrows point to cells shown in the insets with individual far-RFP, RFP and GFP channels. DAPI (blue) marks DNA scale bar is 10μM. **(B-G)** G-TRACE of EC in control young midgut expressing either UAS-LacZ (B, D, F), or UAS-Not-RNAi (C, E, G). Arrows point to cells shown in the insets with individual far-RFP, RFP and GFP channels. **(H)** G-TRACE-based quantification of PPC types (wildtype, PPC* PPC**) observed in control midguts or where Not was targeted ECs. **(I-O)** Confocal microscopy of midguts expressing MyoIA^ts^> G-TRACE system using the indicated antibodies. (B, J, L, N) G-TRACE of EC in young, and (I, K, M, O) old midguts. (P). Quantification of indicated PPCs expressing Odd-Skipped, and Delta similar to experiments shown in L-O.

The phenotypes observed upon acute loss of Non-stop are highly similar to the ones observed in aged midguts (Figure 2I-P; Figure 2 Supplemental Figure 2, 3). G-TRACE analysis of aged ECs established that the aged midgut (5 weeks old) are populated with ECs that are no longer differentiated (PPC**), as well as mis-differentiated progenitors (PPC*). These PPC** no-longer expressed the differentiated LamC, or the transcription factors Pdm1 and Odd-skipped and ectopically expressed the stem cell marker Delta (Figure 2I-O, quantitated in Figure 2P, and Figure 2 supplemental Source data; and Figure 2 Figure Supplemental 3A-D).

As in the case of young ECs lacking Non-stop, aging ECs ectopically expressed the stem cell enhancer M5-4 (Figure 2 Figure Supplemental 2 A, B, I, J). They also exhibited reduced expression of the differentiated Lamin LamC (Figure 2 Figure Supplemental 2C, D, K, L) and ectopic expression of the stem cell-related Lamin, LamDm0 as well as its binding partner Otefin (Ote) in PPC (Figure 2 Figure Supplemental 2E, F, M, N, G, H). At the tissue level, aged ECs also exhibited disorganized distribution of EC-related adhesion molecule Disc large, (Dlg), and reduced expression of MESH and snakeskin (SSK). They ectopically express Armadillo (Drosophila β-catenin), which s expressed on the surface of progenitors in young midguts, all resulting in loss of gut integrity (Figure 2 Figure Supplemental 3E-L; Figure 9O, P). Thus, acute loss of Non-stop in young EC or age-related declines in aged ECs results in EC cells that lose differentiation status and/or mis-differentiate.

**Figure 9:**
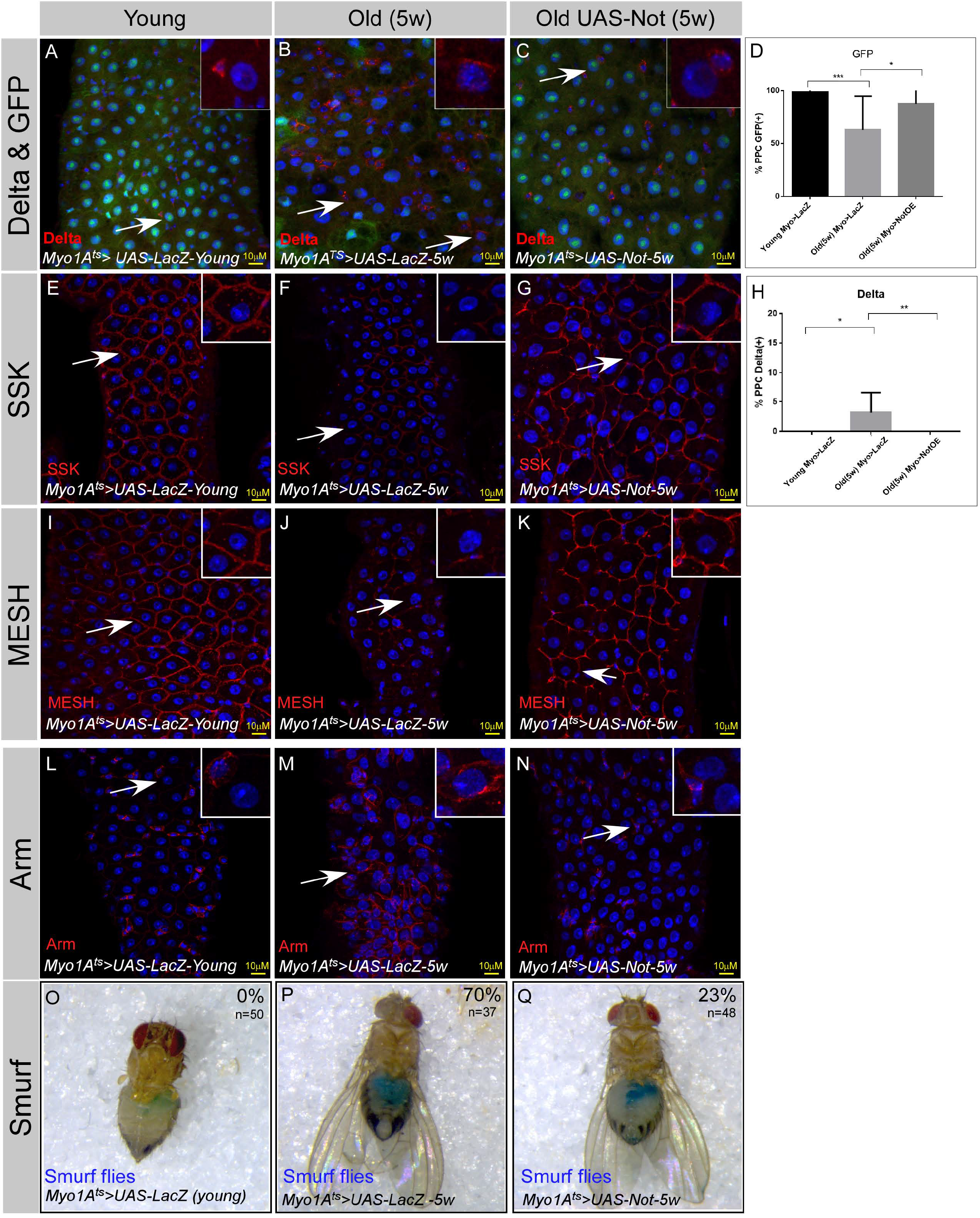
Continues Expression of Not in ECs suppresses aging phenotypes in the midgut. **(A-N)** Confocal images of the midgut tissue using the indicated antibodies and expressing the indicated transgenes in EC using the MyoIA-Gal4/Gal80ts. DAPI marks DNA and scale bar is 10μM. **(A, E, I, L)** midguts derived from 2-4 days old flies (young) **(B, F, J, M)** Midguts derived from 5 weeks old flies expressing control (UAS-LacZ). **(C, G, K, N)** Midguts derived from 5 weeks old flies expressing UAS-Not. DAPI marks DNA (blue), and scale bar is10μM.Arm, Armadillo; SSK Snakeskin. **(D, H)** Quantification of similar experiments shown in (A-C). **** =*P* < 0.0001, \*\*\**P* < 0.001; \*\**P* <0.01; *=*P* <0.1 **(O-P)** Aging impairs gut integrity as evident by the leakage of blue-colored food into the abdomen (smurf assay). Continues expression of Not but not control using the MyoIA-Gal4/Gal80ts for five weeks safeguards gut integrity, n= 48, 38 respectively; *P* < 0.001, **Figure 9; Source data:** Quantification of cell populations described in 8J-L.

### A Non-stop identity complex (NIC) supervises EC identity

Non-stop is the catalytic subunit of a DUB module containing Sgf11, E(y)2 and in some cases Ataxin7 that is part of the SAGA chromatin remodeling complex (Morgan and Wolberger 2017, Mohan 2014, Weake 2008). We therefore tested whether the SAGA complex is required for maintaining EC identity. EC-specific RNAi-mediated reduction of Ataxin7 (part of the DUB module), or GCN5 (the histone acetyl transferase of the SAGA complex) did not result in loss of EC identity (Figure 3, Sup. Figure 1 and not shown). We concluded that the EC identity-related function(s) of Non-stop are independent of SAGA.

Therefore, we biochemically searched for Non-stop-associated proteins that potentially together maintain EC identity. Toward this end, we generated a *Drosophila* S2 cell line stably expressing epitope-tagged Non-stop-2xFLAG-2xHA (Non-stop-FH) under the control of a copper-sulfate-responsive metallothionein promoter. Protein complexes that contained Nonstop were affinity purified using sequential capture of the epitope tags, FLAG, then HA. These complexes were subsequently resolved according to size, using gel filtration chromatography (Figure 3A-C). We used a ubiquitin-AMC de-ubiquitinase activity assay to track enzymatically active Non-stop in the purified fractions (Figure 3B). We found three major peaks of de-ubiquitinase activity. The major activity peak resolved at about 1.8 MDa, together with components of SAGA complex (Group 1). A second peak was resolved centering approximately around 670 kDa (Group 2). A third peak, with the lowest total activity, was detected centering around 75 kDa. The three fractions comprising the center of each peak were combined and constituent proteins identified by mass spectrometry (MudPIT) (Washburn et al., 2001). Group 2 contained e(y)2 and Sgf11but no other SAGA subunits. Remarkably, it also contained members of a known boundary complex that includes Cp190, Nup98, Mod (mdg4) and is known to be part of nuclear complex regulating enhancer-promotor interactions and affecting transcriptional memory (Pascual-Garcia et al 2017). We also noted that Histones H2A and H2B were also detected in both groups 1 and 2, showing the DUB module was co-purifying with known substrates of Non-stop and indicating the DUB was purifying in a physiologically native state (Figure 3C; Zhao et al., 2008).

**Figure 3:**
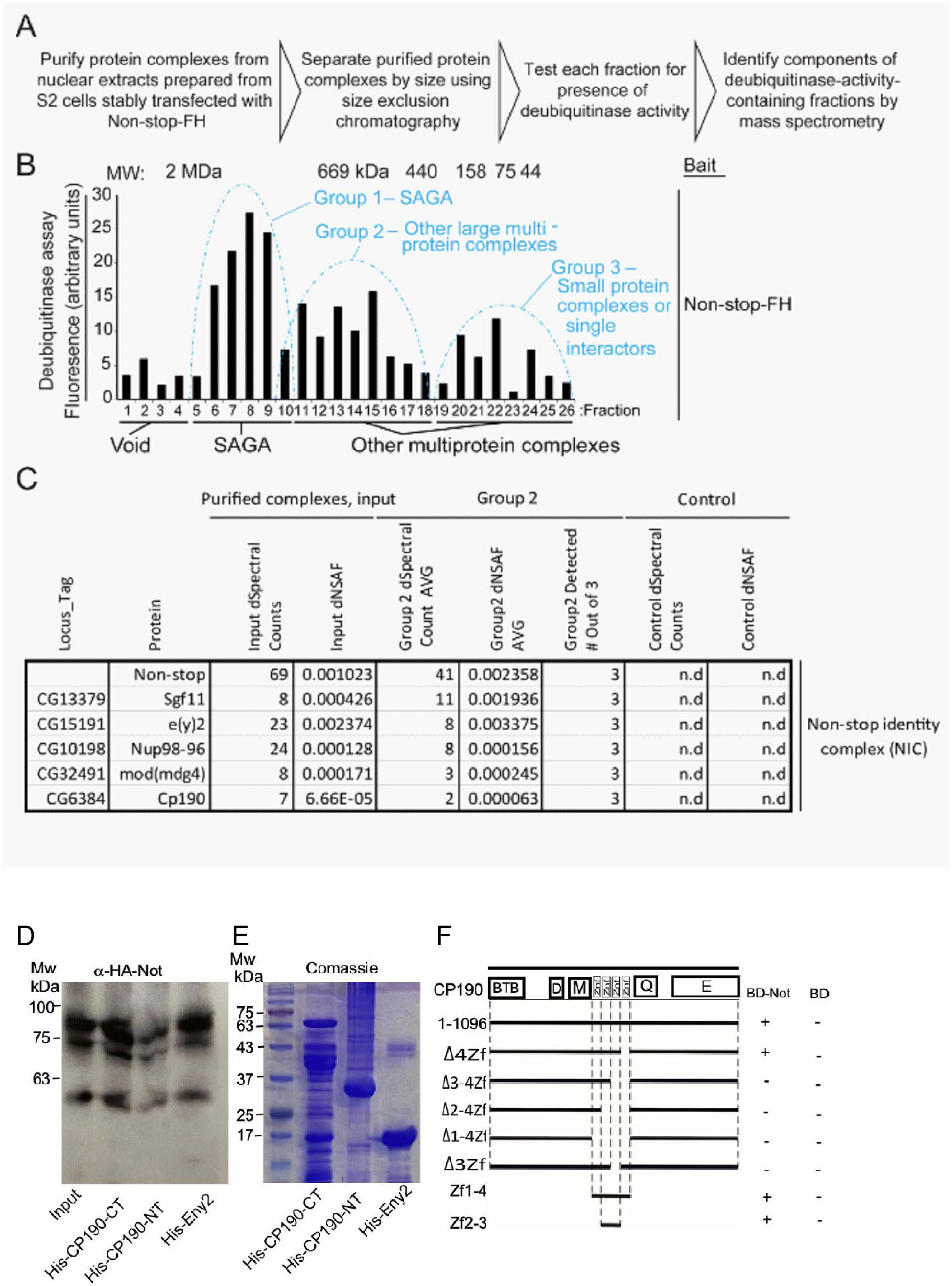
Identification of a Non-stop-identity complex (NIC). **(A-C)** Purification scheme of nuclear Not-associated complexes from *Drosophila* S2 cells. (see text and methods; adopted from Cloud et al 2019) **(B)** Identification of Not-associated isopeptidase activity proteins by immunoprecipitation followed by size fractionation and mass-spectrometry. CP190, Mod (mdg4), Nup96-98, and E(y)2 were all present in Group 2. Not-FH; IP with full length Not FLAG-HA tagged **(C)** Summary of protein complexes isolated identified by mass-spectrometry **(D, E)** Not binds to the C-terminal portion of CP190 and to E(y)2. **(D)** Western-blot of *in vitro* binding between HA-Not derived from S2 cell extract and the indicated bacterially expressed purified His-tagged proteins. 10% input is shown. **(E)** Comassie blue staining of the indicated bacterially expressed His-tagged proteins used in the binding assay in (D). **(F)** Schematic diagram of Y2H interaction assay between CP190 and Non-stop. Different fragments of CP190 were fused to the activation domain (AD) of GAL4 and tested for interaction with Non-stop fused to the DNA-binding domain (BD) of GAL4. Protein domains of full-length CP190 are indicated as boxes, and lines represent the different deletion fragments. Zf denote zinc-fingers; BTB, BTB/POZ domain; D, aspartic acid-rich region; M, microtubule-interacting region; E, acid glutamate-rich region of CP190. The results are summarized in columns on the right (BD-Not and BD alone), with the “+” and signs denotes presence and absence of interaction, respectively.

We mapped the interaction of Non-stop with members of the complex using *in vitro* binding assays and yeast two-hybrid system (Y2H, Figure 3D-F). *In vitro* binding, using S2 cell-derived extract expressing HA-Non-stop and His-tagged proteins, established that Non-stop interacted with its known interaction partner e(y)2 as well as with the C-terminal portion of Cp190 (amino acids (a. a.) 468-1096), but minimally with the N-terminal portion of Cp190 (a.a. 1-524). Additionally, in the Y2H system, Non-stop interacted with full length Cp190. Y2H mapped this interaction to the second and third zinc fingers of Cp190 but not the first or fourth (Figure 3F). Non-stop did not interact with either Nup98 or Mod (mdg4) in similar binding assays (not shown).

We termed this complex NIC (Non-stop identity complex) and hypothesized that if the NIC supervises EC identity, RNAi-mediated elimination of each of its subunits will result in loss of EC identity similar to the loss of Non-stop. Indeed, EC-specific knockdown of all NIC subunits except Sgf11 resulted in loss of identity and inability to maintain expression of the EC gene LamC (Figure 4A-F, Figure 3 Figure Supplemental 1). It also resulted in ectopic expression of Delta (Figure 4G-L). In contrast, loss of Su(Hw), an insulator protein that binds to Mod (mdg4) but was not identified as a Non-stop binding partner, did not result in any detectable phenotype (Figure 3, Figure supplemental 1E).

**Figure 4:**
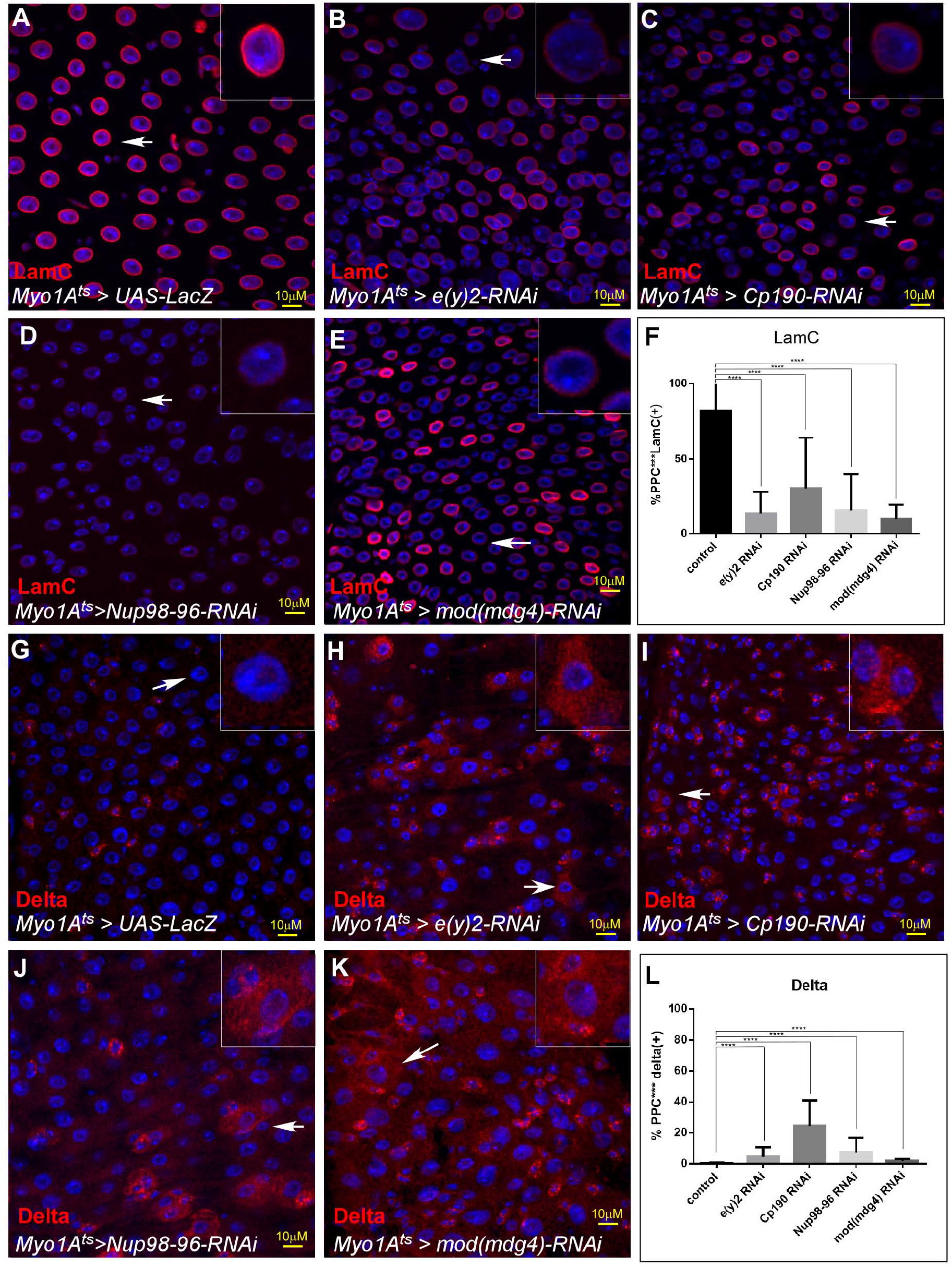
Not identity complex (NIC) regulates EC identity. Confocal images of the midgut tissue using the indicated antibodies; **(A-E)** LamC, **(G-K)** Delta, DAPI marks DNA (blue). The indicated transgenes were expressed in EC using the MyoIA-Gal4/Gal80ts system for fortyeight hours. UAS-LacZ (A, G) UAS-e(y)2-RNAi (B, H); UAS-CP190-RNAi; (C, I). Nup98-96 (D, J) Mod (mdg4) (E, K) White arrows points to cells shown in insets, and scale bar is 10μM. Quantification is shown in (F) for LamC. and (L) for Delta. **Figure 4 Supplemental source data :** Quantification of cell populations described in 4F, 4L.

### Non-stop supervises EC-gene signature and regulates chromatin accessibility

Non-stop is well known to regulate gene expression (Mohan et al. 2014, Li et al. 2017). To elucidate Non-stop-dependent expression signatures, we determined the changes in transcriptional expression using RNA-Seq and its effect(s) on chromatin accessibility by ATAC-seq analyses (Figure 5). We determined the changes in transcription signatures of whole guts upon elimination of Not in ECs using UAS-Not-RNAi and the EC-specific MyoIA-Gal4^ts^. We identified 863 genes with downregulated mRNA expression upon loss of Not in ECs (Table S1 and Figure 5 Figure Supplemental 1D). Of these, 38% (398/1039) were previously identified as EC-related genes (Figure 5A; Korzelius 2014). Metascape analysis unveiled that these shared targets consist of core EC pathways that execute many of the physiological tasks of the gut (Figure 5B; Figure 5 Supplemental Table 1, 2).

**Figure 5:**
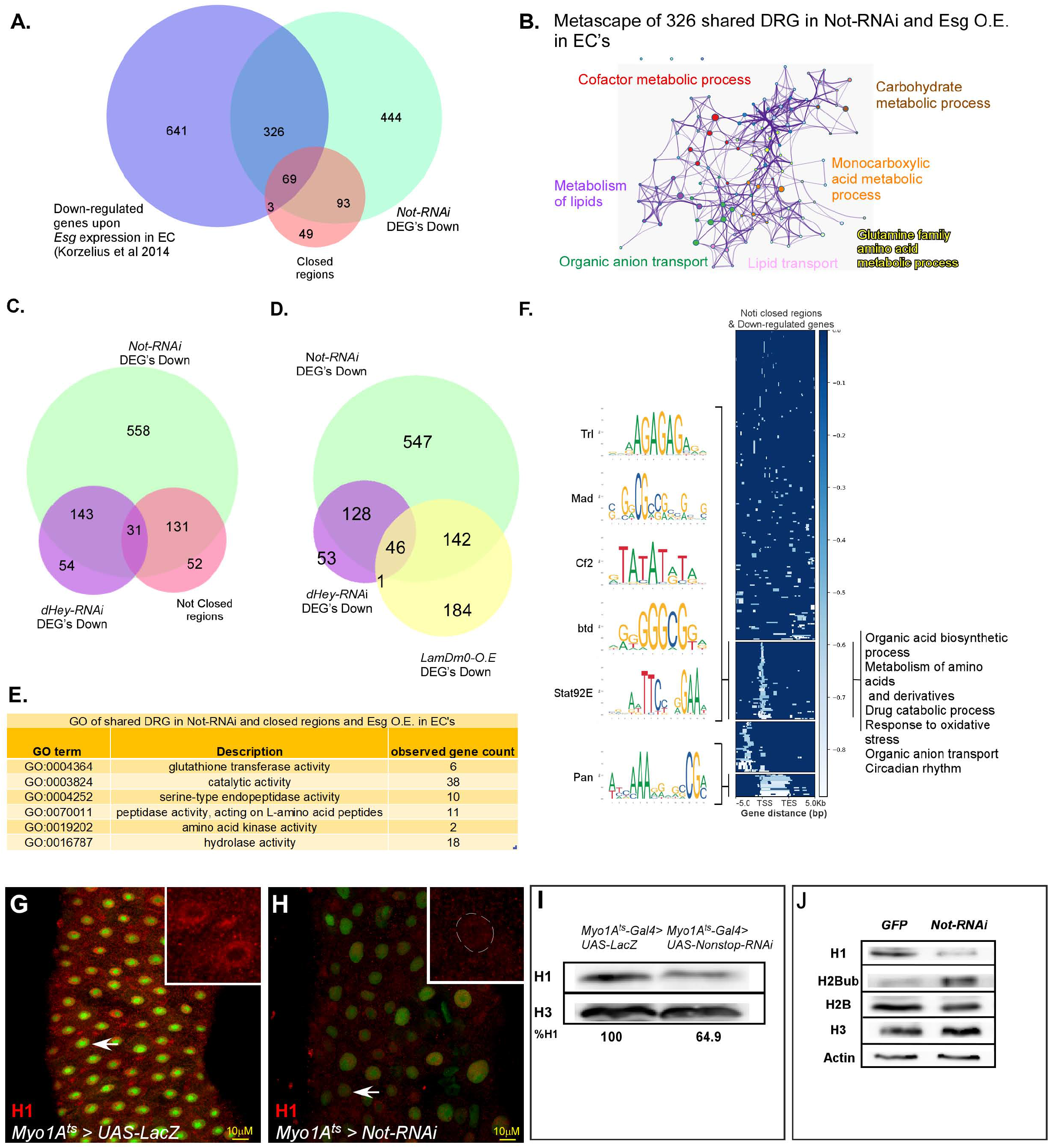
Not regulates EC-gene expression and is required for chromatin accessibility. **(A)** Venn diagram comparing EC-related genes (Blue; Korzelius 2014), genes exhibiting reduced expression upon loss of Not in ECs (Green), and chromatin regions with reduced accessibility upon loss of Not in ECs identified by ATAC-seq (Orange). **(B)** Metascape analysis of Not-down-regulated genes in ECs. **(C)** Venn diagram comparison of genes that exhibit reduced expression upon loss of either Not or Hey in ECs, as well as genes in the vicinity of regions showing reduced accessibility upon loss of Not. **(D)** Venn diagram of genes that exhibit reduced expression upon loss of Not or Hey and of genes with reduced expression upon over expression of LamDm0 in ECs **(E)** GO analysis of genes downregulated by loss of Not in ECs exhibiting reduced accessibility. Observed gene count; number of genes identified from this group in both ATAC-seq and RNA-seq **(F)** Genome-wide alignment and MEME analysis of regions with reduced accessibility in the vicinity of down-regulated genes upon loss of Not in ECs. TSS, transcriptional start site; TES, transcription end site. **(G-H)** Confocal images of the midgut tissue using a-Histone H1 (red), and expressing the indicated transgenes in ECs using the MyoIA-Gal4/Gal80ts system for forty-eight hours, DAPI marks DNA (blue). (G) UAS-LacZ (control) (H,) UAS-Not RNAi. Scale bar is 10μM. **(I, J)** western-blot analysis of the indicated proteins derived from gut extract (I), or S2 *Drosophila* cell extract (J) Histone H3 and Actin serve as loading controls.

We previously identified genes that required Hey for their expression in ECs, and 76% (174/228) of Hey-dependent genes also required Not for expression (Figure 5C).

Moreover, the expression of EC-specific genes was repressed by ectopic expression of the ISC-related lamin, LamDm0, in ECs (Flint-Brodsly 2019). Fifty percent (188/372) of genes that are repressed by expression of LamDm0 in ECs also required Not for their expression, and 46 of these genes were regulated also with Hey (Figure 5D, Figure 5 Figure supplemental 1A-B and see discussion).

In parallel, we examined whether expression of EC-genes involves Non-stop-dependent regulation of chromatin accessibility using ATAC-seq. We identified 214 loci that exhibited reduced chromatin accessibility (“closed”). Of these, 75% (162/214) were located in the range of 0-10Kb vicinity of genes that exhibited reduced expression (Figure 5A; Figure 5 Supplemental Figure 3A, and Supplemental Table S3, S4). GO analysis of these “closed” regions suggested that they belong to genes that maintain the physiological properties of enterocytes (Figure 5E). Alignment of the “closed” chromatin regions showed that they cluster to discrete gene regions (Figure 5F; for EC-related down-regulated genes, and Figure 5, Supplemental Figure 2A for all closed sites, Supplemental Table S5). As shown in Figure 5F, a one cluster was located to the 5’ UTR, a second cluster was at the transcriptional start site (TSS), a third was spanning the coding region, and a fourth was located at the 3’-UTR. MEME analysis revealed that they are statistically significantly enriched in DNA motifs that are known binding sequences of TFs (Figure 5F, Figure 5 Supplemental Figure 2B).

In addition, 565 genes showed upregulation of mRNA expression upon loss of Non-stop, and are related to progenitor fate, cell cycle, and DNA repair (Figure 5, Supplemental 1C). In contrast to numerous closed regions only a small number (~16) regions exhibited increased accessibility upon loss of Non-stop in ECs interestingly many of these genes code of long noncoding RNA (Figure 5 Figure supplemental 2C). Thus, supporting the notion that Non-stop acts primarily to maintain chromatin accessibility in the vicinity of its targets.

We hypothesized that the ectopic expression of these genes may be, at least partially, due to changes in nuclear organization in ECs. In this regard, among the genes that require Non-stop/NIC at the protein level is LamC (Figure 4A-F). LamC is the dominant lamin in ECs that silences the expression of stem cell and non-relevant gene programs in ECs (Flint-Brodsly 2019). For example, PCNA is not expressed in control ECs, but is ectopically expressed in PPC** (ECs that are no longer differentiated; PPC^GFP+RFP-^). This ectopic expression was prevented by co-expression of LamC in ECs where Non-stop was eliminated (Figure 5, Supplemental Figure 1E-G). Moreover, loss of Non-stop in ECs also resulted in a significant decrease in the linker histone H1 that is associated with compacting chromatin and gene silencing (Fyodorov et al. 2018). H1 protein levels were reduced in the nuclear periphery of ECs lacking Non-stop (Figure 5G, H), in gut extracts derived from flies where Non-stop was targeted in ECs, as well as upon knockdown of Non-stop in S2R cells (Figure 5I, J respectively). Thus, the ectopic expression of non-EC programs may be due loss of LamC and H1 proteins and subsequently heterochromatin impairment and dependent silencing. However, since we isolated mRNA from the entire midgut, the source of these upregulated mRNAs may also be from mis-differentiated stem cells (PPC*), as well as from the increase in rapidly dividing progenitor cells.

Comparison of Non-stop RNA-seq data with a genome-wide high-resolution DamID binding map of histone H1 performed in Kc167 *Drosophila* cells (Braunschweig et al., 2009) identified the GAGAGA sequence as the binding sites for the transcription factors Trithorax-related (Trl/GAF), a shared motif for Non-stop-regulated genes also bound by H1. Moreover, GAGAGA sequence was also enriched in of Non-stop closed regions at the TSS of genes requiring Non-stop for expression (Figure 5F).

While Trl/GAF was not identified as part of the NIC in our proteomic purification, it associates in a protein complex containing Nup98, e(y)2 and Mod (mdg4) that regulate gene-expression (Pascual-Garcia et al. 2017). However, targetnig Trl/GAF or the adaptor protein CLAMP protein which bind to the GAGAGA sequence and associates with Cp190 and Mod (mdg4) did not result in loss of EC identity (Not shown; Bag et al. 2019). Therefore, we suggest that NIC is likely functionally and compositionally distinct from the Trl-containing complex.

### Non-stop stabilizes NIC subunits, and Non-stop expression of in aged ECs restores large-scale nuclear organization of ECs and suppresses aging phenotypes

As an isopeptidase, it seemed possible that Non-stop’s ability to maintain expression of EC-related genes stems also from protecting NIC subunits from degradation. Indeed, the protein levels of Cp190, e(y)2, Mod (mdg4), and were reduced upon RNAi-dependent Non-stop elimination in young ECs as observed by immunostaining (Figure 6; Figure 6 Source data). Moreover, Nup98 was no-longer confined to the nuclear envelope but was localized to the nucleus interior in a punctate pattern (Figure 6E, F; Figure 6 Source data).

**Figure 6;.**
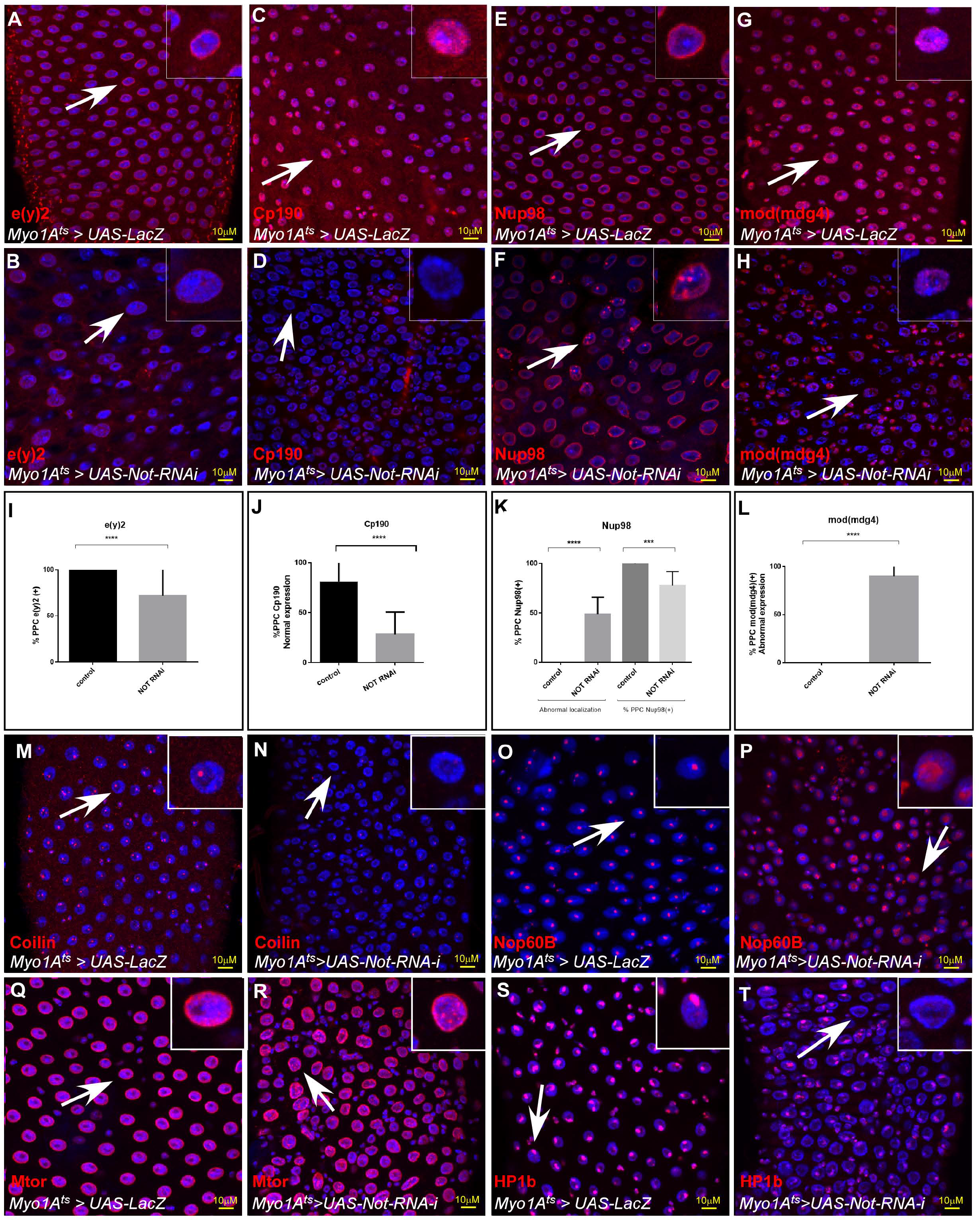
Not maintains the protein level and intranuclear localization the NIC subunits, **(A-H)** Representative confocal images of the midgut tissue using the indicated antibodies (red) and expressing the indicated transgenes in EC using the MyoIA-Ga14/Gal80ts system. UAS-LacZ (A, C, E, G), UAS-Not RNAi (B, D, F, H). DAPI marks DNA (blue), and scale bar is 10μM. White arrows points to cells shown in insets, and scale bar is 10μM. **(I-L)** Quantification of 3 biological experiments is shown **(M-T)** Non-stop regulate large-scale organization of the nucleus. Representative confocal images of the midgut tissue using the indicated antibodies (red) and expressing the indicated transgenes in EC using the MyoIA-Ga14/Gal80ts system. UAS-LacZ (M-P), UAS-Not RNAi (Q-T).

The observed changes in the stability of LamC and NIC subunits encouraged us to examine the larger organization of the nucleus using proteins that are markers for specific intranuclear domains and bodies. Loss of Non-stop in ECs resulted in decline in Coilin, which resides within Cajal bodies, and expansion in the expression of nucleolar Nop60B, a marker of the nucleolus. At the nuclear periphery we noted changes in localization of mTor and subsequent decline in LamC and reduced protein level of HP1b that is associated with heterochromatin and the chromocenter (Figure 6M-T; Figure 2G, H). Moreover, these changes were also observed upon targeting individual NIC subunits (Figure 6, Figure Supplemental 1).

As described above, the physiological relevance of a failure of Non-stop is significant to aging (Rodriguez-Fernandez et al. 2020). The cellular and tissue phenotypes associated with acute loss of Non-stop highly phenocopy aged midguts (Figure 2, Figure supplemental 2, 3). Indeed, a decline in the protein level of Non-stop was observed in aged ECs (Figure 7A, F, P). Moreover, the protein levels of NIC subunits CP190, e(y)2, and Mod (mdg4) were also reduced in aged ECs (compare Figure 7B-E to 7G-J; quantified in Figure 7P-T; Figure 7 source data). Thus, suggesting that a decline in Non-stop protein resulting in a failure to safeguard NIC stability accompanies aging. Therefore, we tested whether preventing the decline in Non-stop protein can protect the loss of NIC. Towards this aim we continuously expressed Non-stop in ECs using UAS-non-stop and the MyoIA>Gal4/Gal80^ts^ system, expressing Non-stop to a level similar to its expression in young ECs as determined by immunofluorescence (Fig 7). Indeed, and consistent with Non-stop’s role as a key stabilizer of NIC expression of Non-stop, but not the control (UAS-LacZ), for five weeks prevented the aged dependent-decline of individual NIC subunits (Figure 7K-O; quantified in Figure 7P-T; Figure 7 source data).

**Figure 7:**
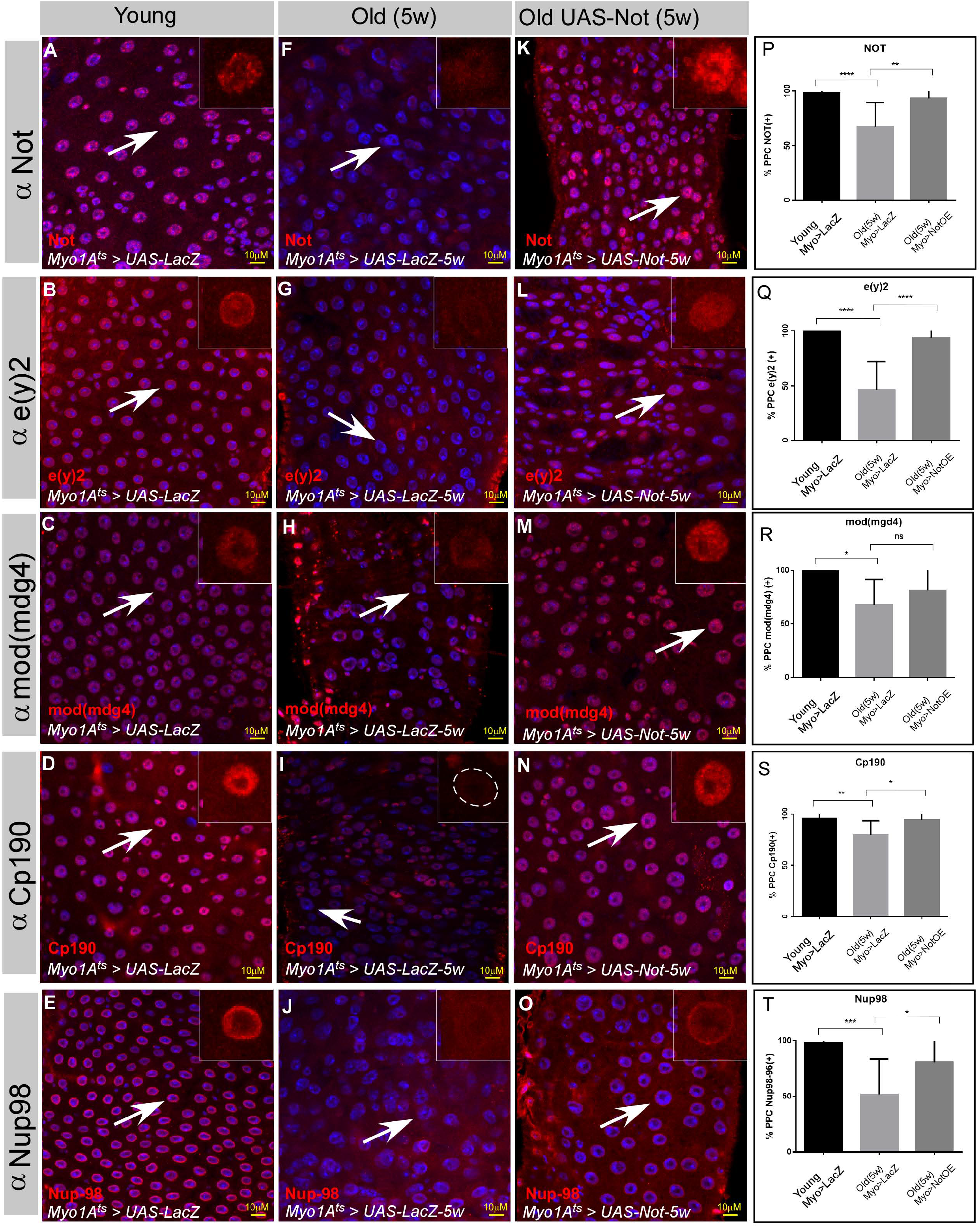
The protein levels of the Not-CP190 complex subunits decline upon aging and is restored upon continues expression of Not in aged ECs. **(A-O)** Representative confocal images of the midgut tissue using the indicated antibodies (red) and expressing the indicated transgenes in EC using the MyoIA-Gal4/Gal80ts system. **(A-E)** Young 4 days old guts, **(F-J)** Five weeks old guts expressing UAS-lacZ. **(K-O)** Five weeks old guts expressing UAS-Not. DAPI marks DNA (blue), and scale bar is 10μM. **(P-T)** Quantification of similar experiments presented in A-O. **** =*P* < 0.0001, \*\*\**P* < 0.001; \*\**P* <0.01; *=*P*<0.1 **Figure 7; source data:** Quantification of cell populations described in 7P-T.

We further examined whether maintaining Non-stop protein levels attenuates the aging of the midgut using the above system. Aging is associated with distorted nuclear organization of ECs (Figure 8, Figure 2, Figure 2 supplemental Figure 2). These changes include re-organization of the nuclear periphery, including a reduced level of LamC and histone H1 as well as redistribution of mTOR (Figure 8A-I). Aging is also associated with ectopic expression of LamDm0, and re-localization to the nuclear periphery of LamDm0 binding partner, Otefin (Ote) (Figure 8J, K, H, and Figure 8 supplemental Figure 1). Changes are also observed in the nucleus interior, involving the nucleolus and Cajal bodies as observed by the expansion of the nucleolar protein Nop60B and Coilin, which are resident proteins in these sub-nuclear bodies (Figure 8M-O; Figure 8 supplemental Figure 1). Consistent with Non-stop as a key identity supervisor relevant to aging, continuous expression of Non-stop for five weeks suppressed the above age-related changes in nuclear organization. Non-stop expression greatly restored LamC, histone H1 levels, localization of Mtor and suppressed the ectopic expression of LamDm0 and Ote, as well as restored the large-scale organization of the aged EC nucleus (Figure 8 C, F, I, L, O and Figure 8 supplemental Figure 1).

**Figure 8:**
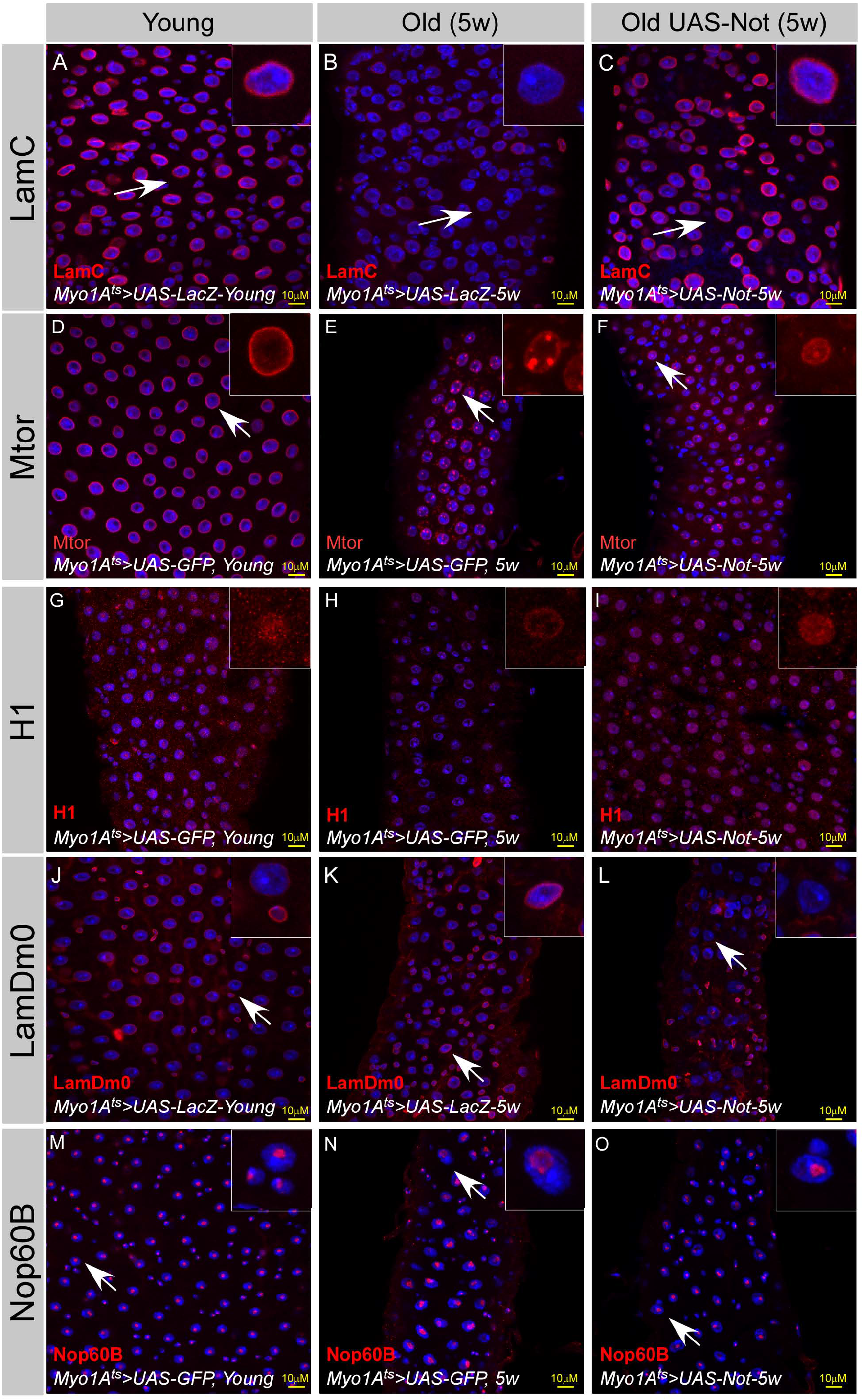
Expression of Non-stop restore large-scale organization of aged ECs: **(A-O)** Confocal images of the midgut tissue using the indicated antibodies and expressing the indicated transgenes in EC using the MyoIA-Gal4/Gal80ts. DAPI marks DNA, and scale bar is 10μM. (A, D, G, J, M) Young Guts expressing UAS-LacZ. (B, E, H, K, N) Five weeks old guts expressing control (UAS-GFP). (C, F, I, L, O) Five weeks old guts expressing UAS-Non-stop. (A-C) a-LamC; (D-F) Mtor; (G-I) a-Histone H; (J-L) a-Lam1Dm0; (M-O) a-Nop60B..

We further tested whether expression of Non-stop in wild-type ECs is capable of attenuating age-related changes in the gut epithelia. Indeed, expression of Non-stop suppressed classical characteristics of the aged midgut. For example, Non-stop expression maintained the expression of the EC marker Myo>GFP and suppressed the ectopic expression of the ISC marker Delta PPCs (Figure 9A-D, H). Continuous expression of Non-stop in wildtype ECs for five weeks restored the expression of EC-related junctional proteins SSK and MESH, and suppressed the ectopic expression of Arm in PPCs (Figures 9E-G, and 9I-N). To test whether Non-stop maintenance affected the entire midgut at the organ level we tested for overall gut integrity using the Smurf assay. We observed that continuous expression of Non-stop in ECs greatly prevented the extensive leakage of blue-colored food observed in five weeks old aged animals restoring gut integrity (Figure 9O-Q). Thus, Non-stop is required for expression of EC-gene programs, stabilizes NIC subunits in the adult, and together with NIC regulates large-scale organization of the differentiated nucleus, safeguarding EC identity and protecting from premature aging.

## Discussion

### An *in vivo* screen identifies regulators that maintain the differentiated state

We performed an identity screen focused on conserved enzymes within the ubiquitin and ubiquitin-like pathways using the midgut tissue as a model system. Based on cell-specific secondary tests, we identified three categories of supervisors; 1. EC-specific identity regulators 2. Genes that are required for differentiated cell identity (both EEs and ECs) 3. Genes that are required for identity of all cell types (general identity regulators). Of specific interest were a group of genes (CG1490; CG2926; CG4080) whose elimination in EEs resulted in a loss of EC, but not EE, identity, acting as inductive identity regulators. Likely their effect on ECs involves cell~cell communication via diffusible factors. The latter may be stochastic, or may be mediated by microtubule-based nanotubes as in the crosstalk between the hub cell (part of the stem cell niche) and stem cells in the *Drosophila* testis (Inaba et al. 2015).

The observation that Ub/UbL-related genes protect the differentiated identity is conserved across species. Screens in mammalian systems identified enzymes within the SUMO and ubiquitin pathways acting as a barrier against forced reprogramming of differentiated cell. Among these genes were Ubc9, the sole SUMO conjugating enzyme, that was also identified in our screen and the isopeptidase Psmd14 (Cheloufi et al. 2015; Buckley 2012). In addition to Non-stop, screen identified the iso-isopeptidase UTO6-like (CG7857), Usp7, and Rpn11 as regulators of EC identity. Rpn11 is part of the lid particle of the 26S proteasome, involved in deubiquitinating proteins undergoing proteasomal degradation (Greene et al. 2020). We also identified the core particle proteasome subunit Pros-a6 (CG4904) as a bona-fide identity supervisor but not other proteasome subunits (that were also screened). Thus, it is not clear whether it represents a unique proteasome-independent function of the Pro-a6 subunit or whether Pro-a6 is a limiting subunit for proteasome biogenesis and activity in ECs.

Identity supervision is intimately involved in cancer, and genes regulating identity are likely to serve as a barrier to tumorigenesis and tumor suppressors. The human ortholog of Non-stop /USP22 has mixed oncogenic and tumor-suppressive functions (Jeusset et al. 2017). Relevant to our study is the observation that USP22 has tumor suppressive functions in colon cancer by reducing mTor activity (Kosinsky et al. 2020). Along this line it is interesting to note that 10/17 of human orthologs to genes discovered in our screen are either mutated or silenced in cancer. Thus, future studies of these human orthologs may identify potent tumor suppressors in cancer.

### Crosstalk between identity supervisors

Both Non-stop and the transcription factor Hey are bona-fide regulators of EC identity required for the expression of EC-related genes. We found a significant number of EC-related genes that required both Non-stop and Hey for their expression, suggesting that Hey and Not may co-regulate these genes. However, functional and epistatic tests suggest that Hey also acts upstream or in additional pathways to Non-stop. Hey binds to enhancers in lamin genes repressing the expression of the ISC-related lamin LamDm0 and enhances the expression of LamC. In contrast, Non-stop does not regulate the accessibility or expression of either LamDm0 or LamC at mRNA level. However, Non-stop is required for maintaining the levels of LamC protein. Therefore, loss of Non-stop result in a decline in LamC but not in the ectopic expression of LamDm0, which is observed upon acute loss of Hey or aging.

This discrepancy may be due to the presence of Hey on its repressed targets in young ECs where Non-stop is targeted, and directly repressing their expression as maybe in the case of LamDm0. Moreover, EC-specific expression of Non-stop did not suppress the phenotypes associated with acute loss of Hey in young ECs further supporting for Hey-dependent, but Non-stop independent functions. However, the ECs-specific expression of either Non-stop or Hey in aging midguts restores expression of LamC and repressed ectopic LamDm0 expression.

### Potential function of Non-stop and the NIC

A possible function of the NIC may be the recruitment of the H3K4me2/3 COMPASS methylases to catalyze H3K9 di- and trimethylation at enhancers and promotors, which are fundamental for gene activation (Shilatifard 2012; Sze and Shilatifard 2016). One prominent phenotype of loss of Non-stop was the mislocalization of Nup98 from the nuclear periphery to intranuclear punctate pattern. Nup98 was shown to recruit Set1~COMPASS to enhance histone H3K4me2-3 methylations in hematopoietic progenitors (Frank et al 2017). Thus, NIC may be required for recruitment of COMPASS and enhancing transcriptional memory promoting the transcription of EC-related genes.

H3K4 methylation and gene activation also require a ubiquitination and de-ubiquitination cycle catalyzed by the Bre1 ubiquitin ligase, and the de-ubiquitinase Non-stop/USP22 (Lee et al. 2007; Nakanishi et al. 2009). It is possible that with respect to EC-related genes, activity of Bre1 or Non-stop/NIC has a direct role in gene transcription in a similar fashion.

Moreover, we found that Non-stop is required for the stability of NIC, possibly by deubiquitinating NIC subunits *in vivo,* and that this stabilization is relevant in the context of physiological. However, the stabilization of NIC subunits maybe a more indirect role of Nonstop and mediated via LamC. We noticed that LamC expression partially restored the protein levels of NIC subunits and their intranuclear localization potentially by serving as a scaffold for NIC at the nuclear periphery. Thus, Non-stop may function at two levels; One is a direct role in transcription within NIC promoting de-ubiquitination of H2Bub while a second function is the stabilization of identity supervisors including NIC subunits and LamC.

### Not LLPs, and pre-mature aging

Changes in large-scale nuclear organization are hallmarks of aging (Zhang W 2020). Expression of identity supervisors can prevent age-related distortion of the nucleus EC identity and protect overall the epithelial tissue (This work and flint Brodsly et al 2019). However, to accomplish this, Non-stop or Hey were continuously expressed in ECs and temporal expression of Hey or Non-stop in already aged ECs was not sufficient to suppress aging phenotypes. Thus, if the levels of identity supervisors are kept at youthful levels, they can continue to maintain cell identity and prevent signs of aging, effectively keeping the gut organization and structure similar to young tissue (Kenyon 2010).

Furthermore, it is not clear how expression of a single regulator like Non-stop has an extensive impact on the entire nucleus. Recent studies suggest that Non-stop functions in additional multiprotein complexes that may regulate large-scale cellular organization. For example, Nonstop is part of an Arp2/3 and WAVE regulatory (WRC) actin-cytoskeleton organization complex where it deubiquitinates the subunit SCAR (Cloud et al. 2019). In this regard, a nuclear actin organizing complex, WASH, interacted with nuclear Lamin and was required for large scale nuclear organization (Varbooon et al. 2015). Thus, it is tempting to suggest that such complexes are required to maintain cell identity, and that subunits within these complexes are deubiquitinated by Non-stop.

Recent studies suggest that the organization of the nucleus is mediated by the biophysical properties of the nuclear protein milieu and interaction with macro-molecules such as chromatin and formation of local condensates (Strom and Brangwynne 2019; Yoshizawa 2020). These biophysical forces including liquid-liquid phase-separation (LLPS), are critical for compartmentalization of the nucleus, heterochromatin and euchromatin formation, establishment of transcription factories and intranuclear bodies. Thus, Non-stop/NIC activity may be critical for safeguarding the stability of proteins that their local concentration is critical for the self-organization and compartmentalization of the differentiated nucleus.

In this regard many nuclear proteins are extremely long-lived proteins (LLPs) among them are nuclear pore complex proteins (NPCs) and core histones (Toyama et al. 2013; Toyama 2019). The extended stability of LLPs may originate from intrinsic properties of LLPs, or due to sequestration and evading degradation. However, increased stability may be also actively maintained by constitutive de-ubiquitination. Indeed, post-transcriptional modification by ubiquitin and SUMO were shown to regulate lamin stability and their intranuclear localization (Blank M. 2020). Specifically, type-A lamin and its splice variant Progerin, the cause of Hutchinson Gilford progeria syndrome (HGPS), a premature aging syndrome, are degraded by the HECT-type E3 ligase Smurf2 via ubiquitin-dependent autophagy (Borroni et al. 2018). The elimination of Progerin by expression of Smurf2 in HGSP-fibroblasts reduced the deformation observed in these cells. Thus, it is possible that enhancing Progerin degradation by inhibiting the human ortholog of Non-stop, USP22, will restore nuclear architecture, and suppress the premature aging phenotypes observed in HGPS cells.

## Supporting information

S1 - Screen Results

S2 - Proteomic analysis

S3 - RNA-seq of Not-regulated genes.

S4 - ATAC-seq profiling

S5 - Gene clustering of Non-stop closed regions

## Authors’ contributions

NE, LI, ELB, OM RM PV, TD and AO designed and preformed experiments, WW and NE preformed genomic and bioinformatics analyses. All authors analyzed data. ELB, RM, TD, and AO wrote the paper.

## Acknowledgments

We are grateful for mass spectrometry done by Skylar Martin-Brown, Laurence Florens, and Michael P Washburn at the Stowers Institute. We would like to thank Sarah Bray, Adi Salzberg, Bruce Edgar, Jeff Reinitz, Lori Wallrath, Pamela Geyer, Yossi Greenbaum, Lorry Pile, Erika Matunis, Díaz-Benjumea, Mikio Furuse Bas Van-Steensel, Moshe Oren, Joseph Gall, the Bloomington, VDRC, and NIG-FLY *Drosophila* stock centers for sharing antibodies, fly lines, reagents, and data.

This research was supported by: The School of Biological and Chemical Sciences, UMKC; University of Missouri Research Board; UMKC SEARCH, UMKC SUROP scholars programs, and NIH Academic Development Via Applied and Cutting Edge Research (ADVANCER) program; NIGMS grant 5R35GM118068 to RM. Washington University School of Medicine / St. Louis Children’s Hospital Children’s Discovery Institute, MC-II-2014-363 to TD. Russian Science Foundation 19-74-30026 to PG, and the Israel Science Foundation (ISF) (Grants 739/15, 318/20), and by the Flinkman Marandi Family cancer research grant to AO.

---

**Figure 1; Figure Supplemental 1:**
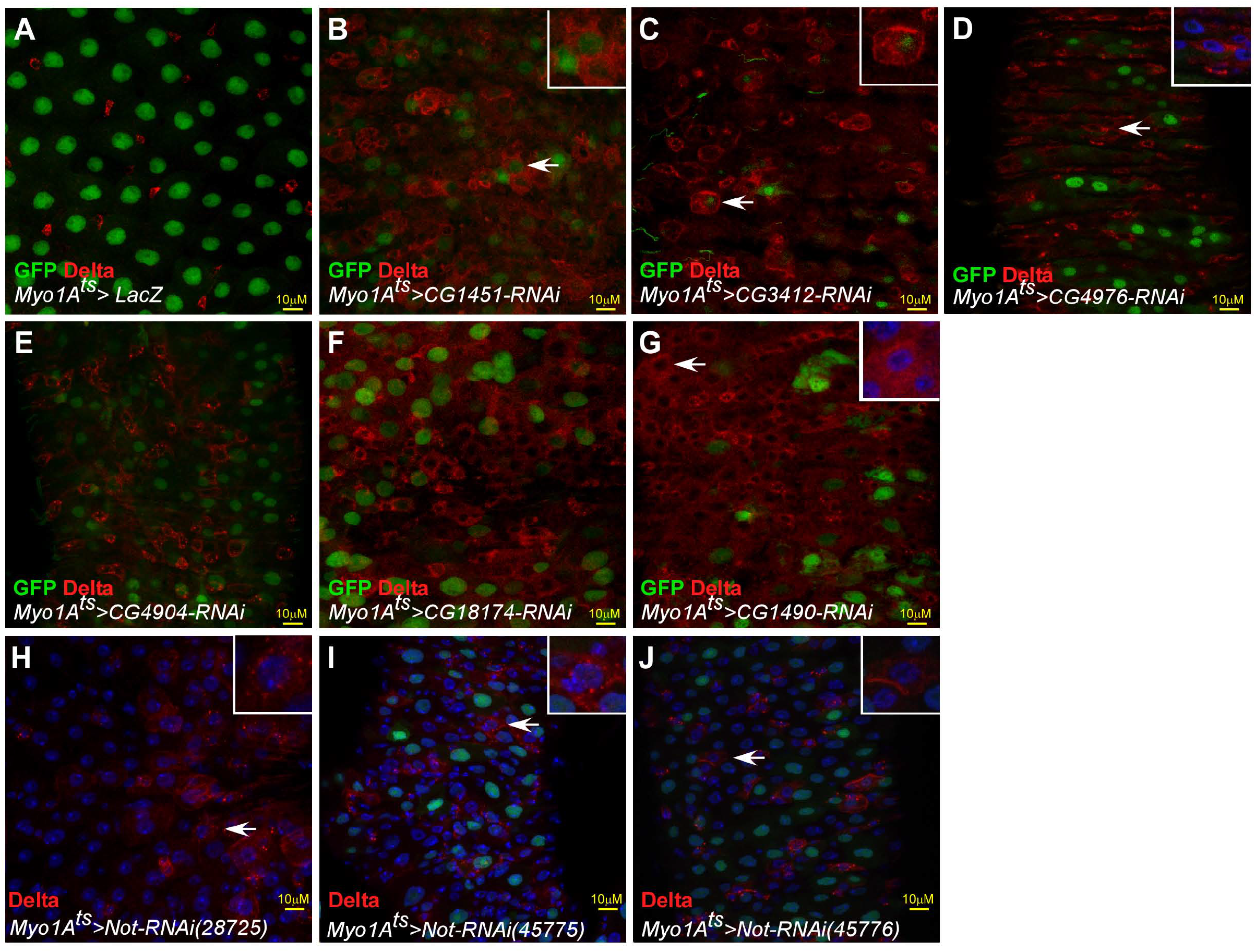
Examples of positive hits of the Ub/UbL screen: **(A-K)** Confocal images of the midgut tissue and the indicated transgenes expressed in ECs using the MyoIA-Gal4/Gal80ts. White arrows indicate cells shown in insets. Scale bar is 10μM MyoIA>UAS-GPF marks fully differentiated ECs, Delta is shown in red, and DAPI (blue) marks DNA. **(A-G)** Examples of positive hits from the screen. **(H-J)** Loss of in ECs Non-stop using three independent UAS-RNAi transgenic lines results in loss of EC identity.

**Figure 1 Figure Supplemental 2:**
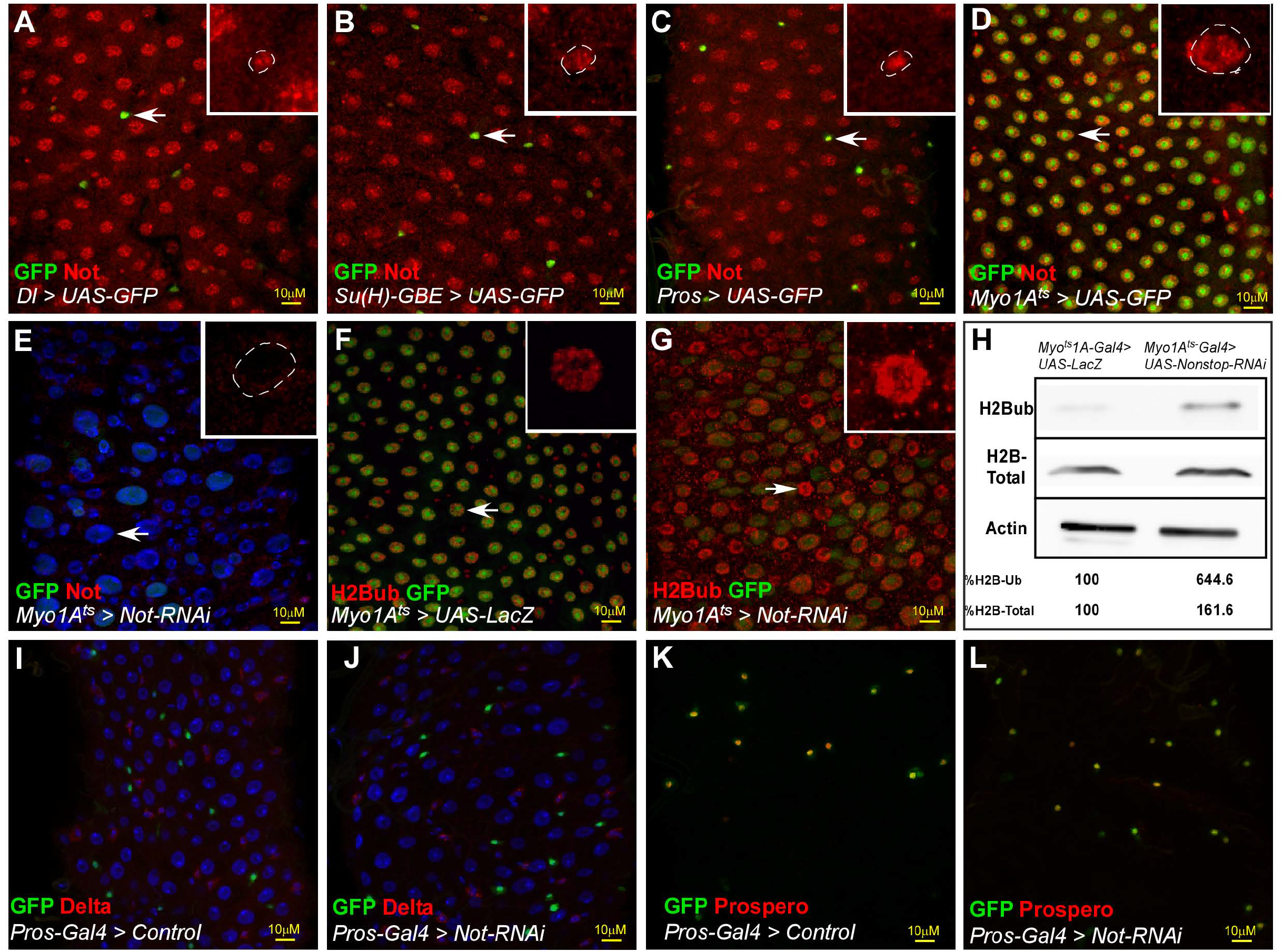
Characterization of Non-stop in midgut cells. **(A-G)** Confocal images of the midgut tissue and the indicated transgenes expressed in EC using the MyoIA-Gal4/Gal80ts. (A-D) Not is expressed in all mid gut cells. Expression of endogenous Not protein (red) was tested relative to the expression of UAS-GFP that was expressed under the cell-specific GAL4 drivers: Dl>GAL4 (ISC); Su(H)>Gal4 (EBs); Prospero Gal4 (EE’s) and MyoIA>Gal4 (ECs). Arrow point to cells shown in insets. (**E)** Expression of Not upon activating UAS-Not-RNAi in ECs. **(F, G)** Level of H2Bub (red) in midguts expressing the control (F), or UAS-Not-RNAi (G) in ECs using the MyoIA>Gal4, UAS-GFP system. **(H)** Western-blot analysis of H2Bub and H2B in midgut derived extracts of the indicated genotypes. Actin serves as a loading control. **(I-L)** Expression of Delta (I, J, red) or Prospero (K, L, red) in midguts expressing control (I, K) or the UAS-Not RNAi (J, L) in EE using *prospero*>GAL4^ts^, UAS-GFP system. Scale bar is 10μM

**Figure 1; Supplemental Table 1:** Summary of the screen Ub/UbL screen results.

**Figure 1; Source data:** Quantification of data presented in Figure 1G, H.

**Figure 2; Figure Supplemental 1:**
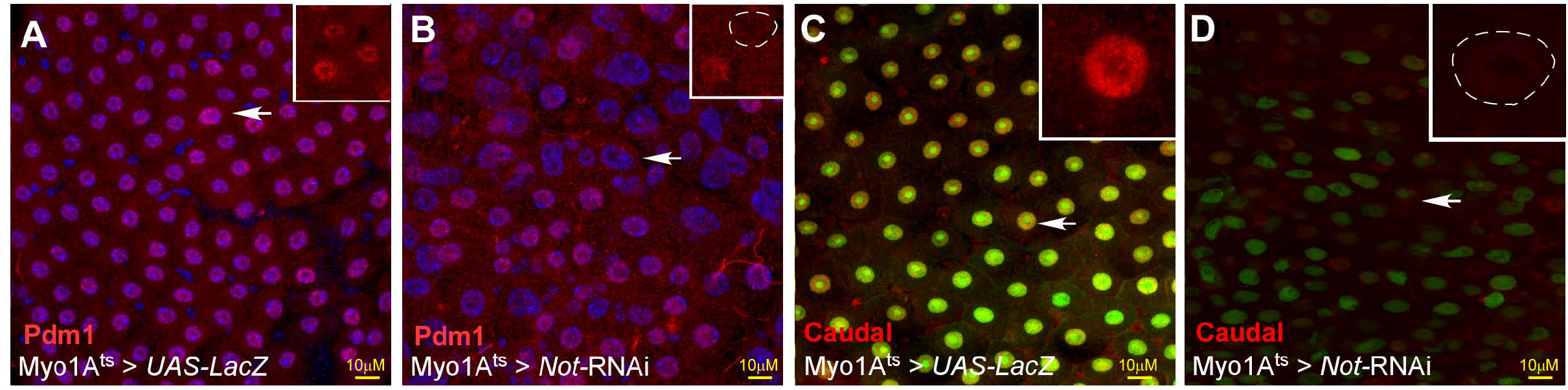
Non-stop is required for expression of EC transcription factors Caudal and Pdm1. **(A-D)** Confocal images of the midgut tissue and the indicated transgenes expressed in EC using the MyoIA-Gal4/Gal80ts. (A, B) anti-Pdm1 (C, D) anti-Caudal Arrows point to cells shown in the inset. Scale bar is 10μM

**Figure 2; Figure Supplemental 2:**
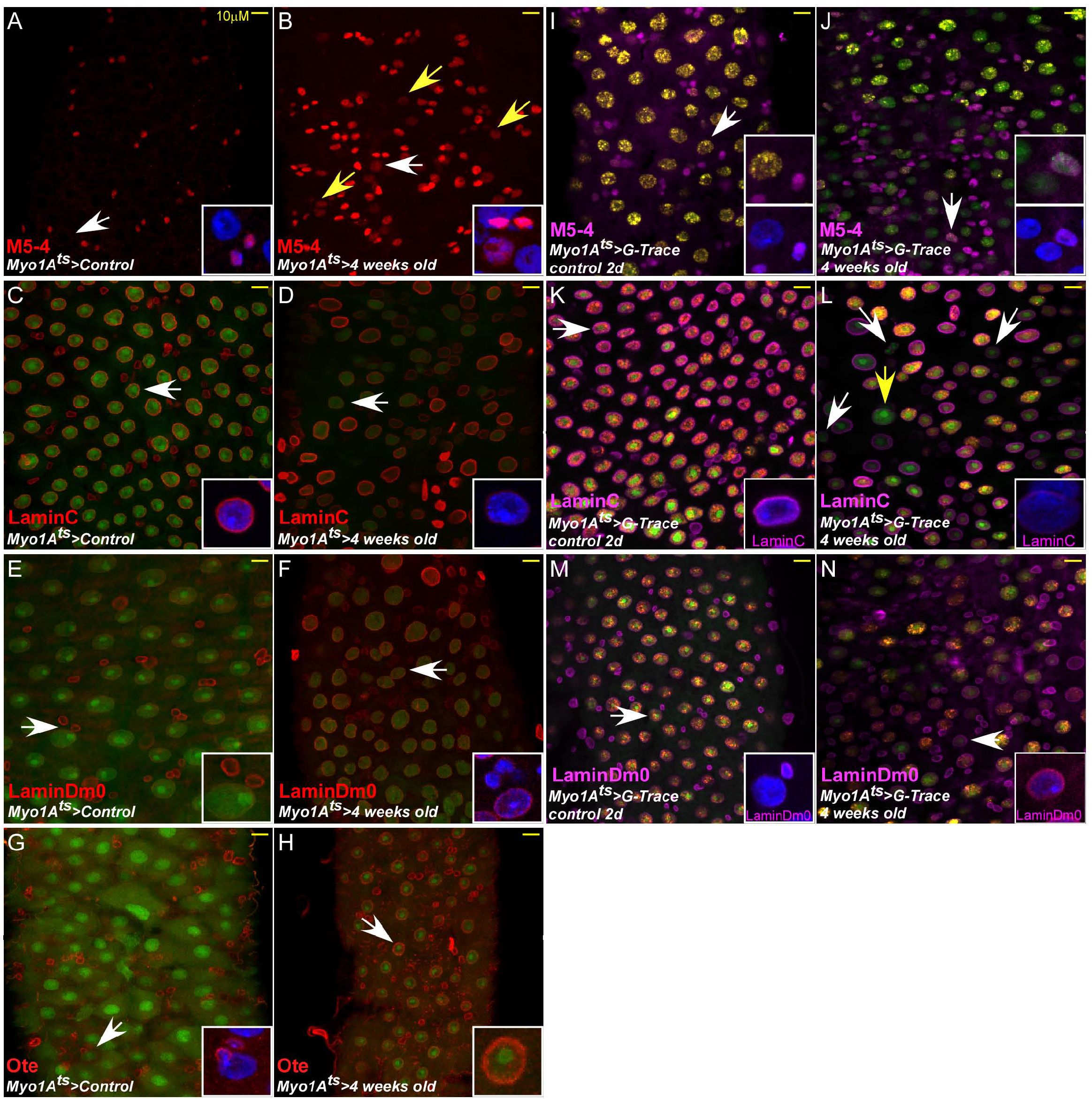
Miss-regulation of enhancers activity and nuclear Lamins in aged enterocytes. **(A-N)** Confocal images with the indicated antibodies of adult *Drosophila* midgut epithelium expressing the indicated transgenes, DAPI marks DNA, and scale bar is 10μm. **(A-H)** transgenic lines expressing UAS-GFP under the control of the EC-specific promoter MyoIA-GAL4/Gal80^ts^ system. **(I-P)** Transgenic lines expressing G-TRACE system under the control of the enterocyte specific promoter MyoIA^ts^ system. The expression of M5-4::LacZ stem cells enhancer of *esg* gene is shown in A, B, I, and J. The protein level and distribution of the indicated protein is shown; LamC expression (C, D, K, L). Lamin Dm0 (LamDm0), E, F, M, N., and Otefin (Ote) (G, H, O, P). Otefin (Ote) protein level in control young ECs or old (G, H).

**Figure 2; Figure Supplemental 3:**
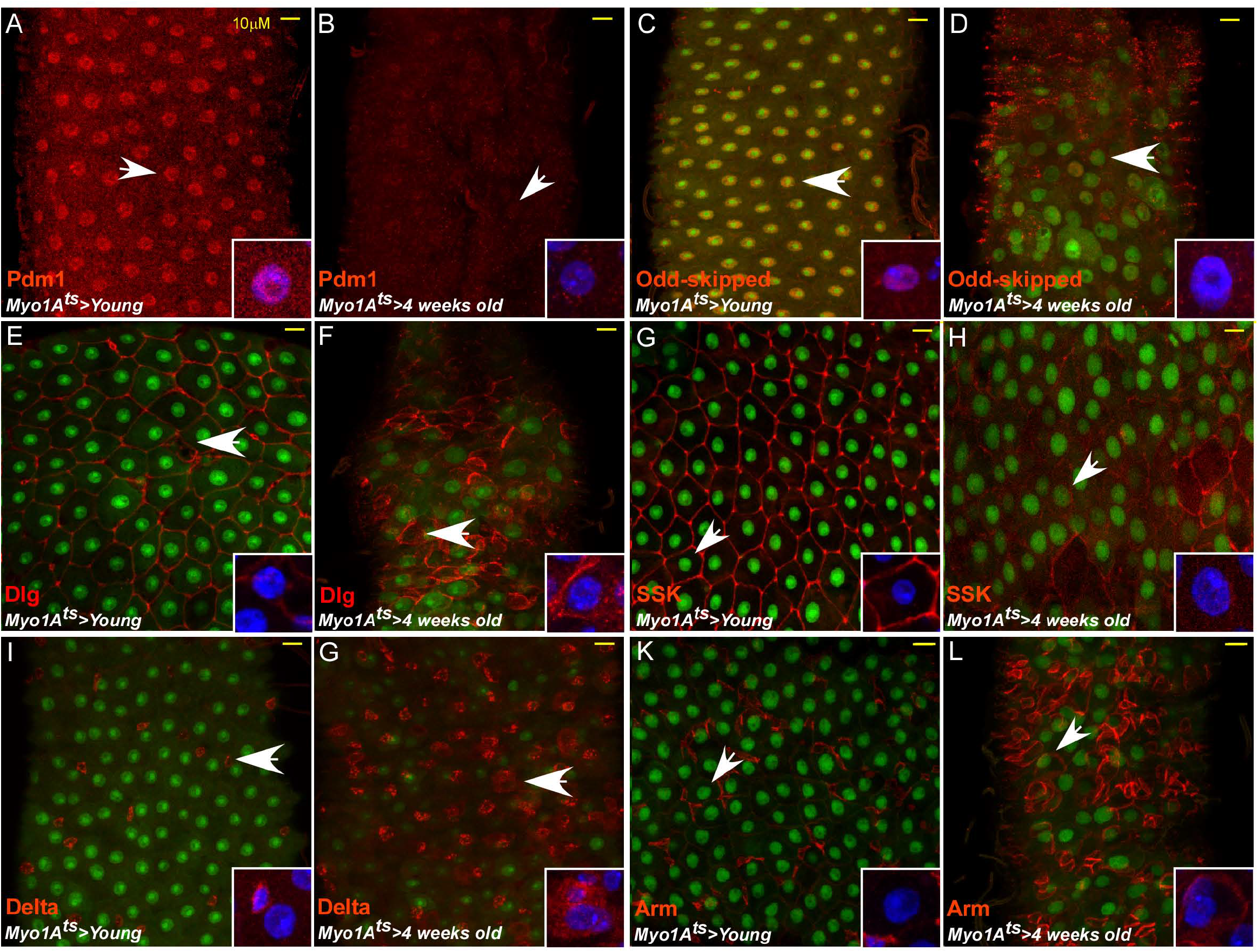
Hallmarks of aging in the *Drosophila* midgut. **(A-L** Confocal images of indirect immunofluorescence staining with the indicated antibodies of adult *Drosophila* midgut intestinal epithelium expressing termed MyoIA^ts^. Scale bar is 10μm. (A, C, E, G, I, K): Young mid-guts (four days old adults). (B, D, F, H, J, L) Four weeks old guts. DAPI marks DNA and arrows indicates cells shown in the insets. SSK; Snake Skin; Arm, Armadillo; Dlg, Disc large. **Figure 2 Source data file:** Quantification data for Figures 2H, 2P,

**Figure 3; Figure Supplemental 1:**
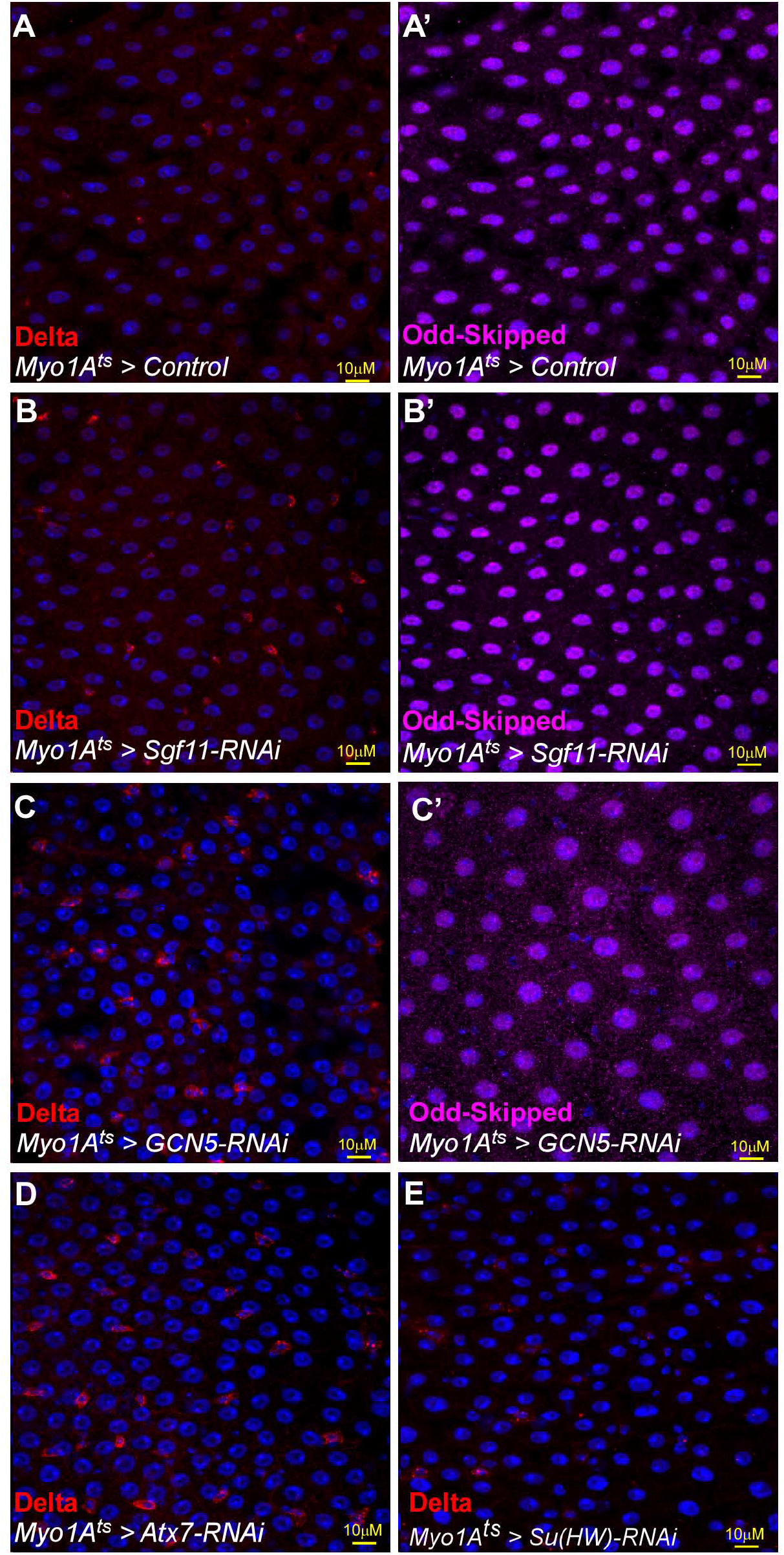
Loss of SAGA subunits and Su(Hw) does-not affect EC identity. **(A-E)** Confocal images of the midgut tissue and the indicated transgenes expressed in EC using the MyoIA-Ga14/Gal80ts. (A-E) anti-Delta (A’-C’) anti-Odd-skipped. (A, A’) UAS-LacZ, (B, B’) UAS-Sgf11-RNAi; (C, C’) UAS GCN5-RNAi, (D) UAS-Atx7 RNAi (E) UAS-Su(Hw)-RNAi. DAPI marks DNA, Scale bar is 10μM.

**Figure 5; Figure Supplemental 1:**
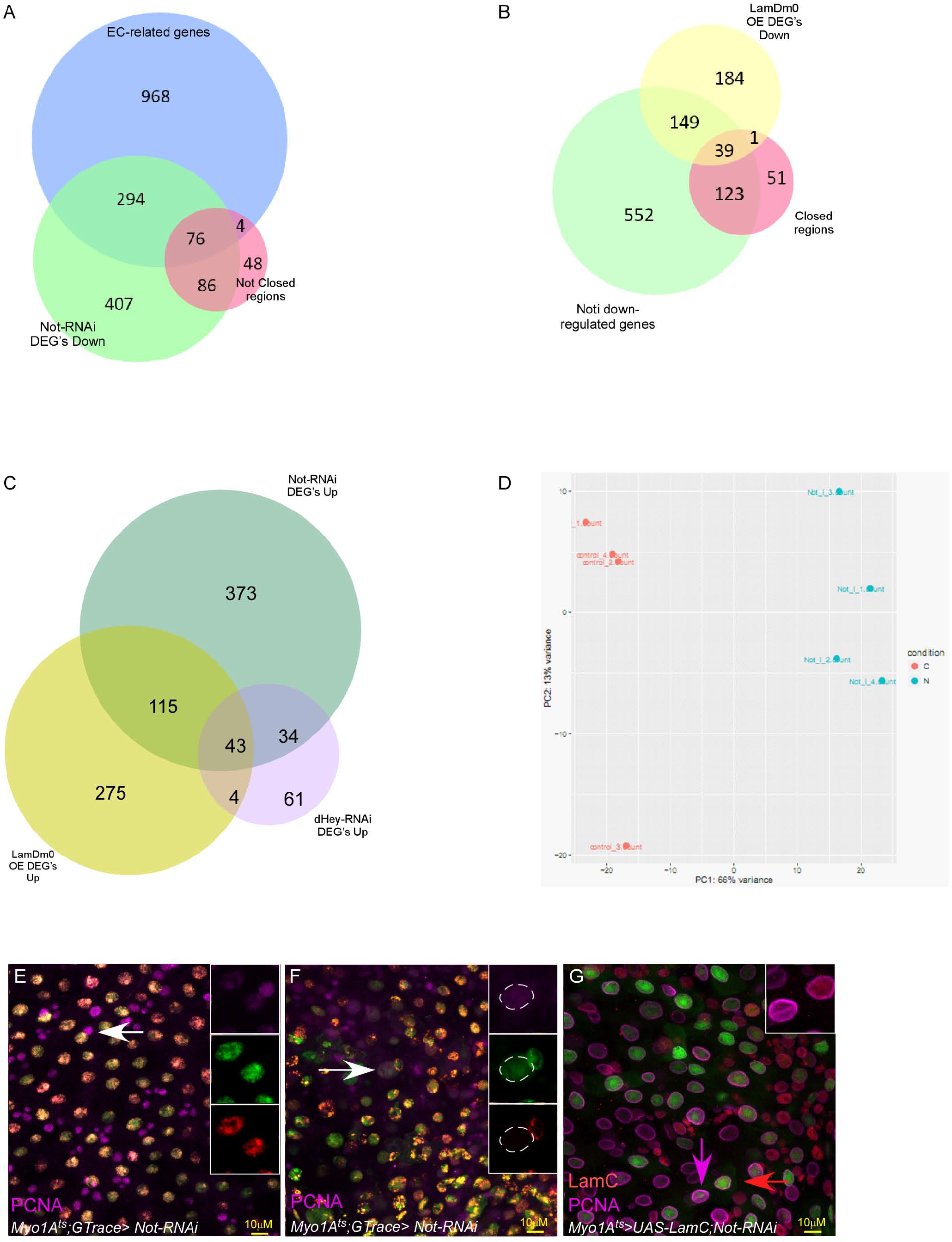
Analysis of Not-related RNA-seq and ATAC-seq. **(A)** Venn diagram comparing genes exhibiting enriched expression in differentiated gut cells (Blue), genes exhibiting reduced expression upon loss of Not in ECs (Green), and chromatin regions with reduced accessibility upon loss of Not in ECs identified by ATAC-seq (Orange). Venn diagram comparison of genes that exhibit reduced expression upon loss of either Not or expression of LamDm0 ECs, and EC-expressed genes **(B)** Venn diagram comparison of genes that exhibit reduced expression or accessibility upon loss of Not, and genes with reduced expression upon over expression of LamDm0. **(C)** Venn diagram comparison of genes that exhibit upregulation in expression upon loss of either Not or Hey in ECs or over-expression of LamDm0 in ECs. **(D).** Three principle components analysis of RNA-seq. **(E-G) Expression of LamC suppresses the ectopic expression of PNCA in EC that are no longer differentiated (PCC**).** Confocal images of the midgut tissue using the indicated antibodies. (E-F) G-TRACE analysis; (E) Control ECs (expressing UAS-LacZ) do not express PCNA, and are both RFP^(+)^ GFP^(+)^. (F) PPC** are GFP^(+)^ and RPF^(-)^ (PPC **) and express PCNA (purple). Arrow points to cells shown in the insets (individual channels). **(G)** ECs where Non-stop was eliminated using the MyoIA-Ga14/Gal80ts system for forty-eight hours ectopically express PCNA (purple), but not in cells that co-express UAS-LamC. Example of two cells is shown; the purple and red arrows point to the cells shown in the inset; The left cell exhibits high level of LamC (red) and low level of PCNA (purple), and the right cell exhibit low level of LamC and high level of PCNA.

**Figure 5; Figure supplemental 2:**
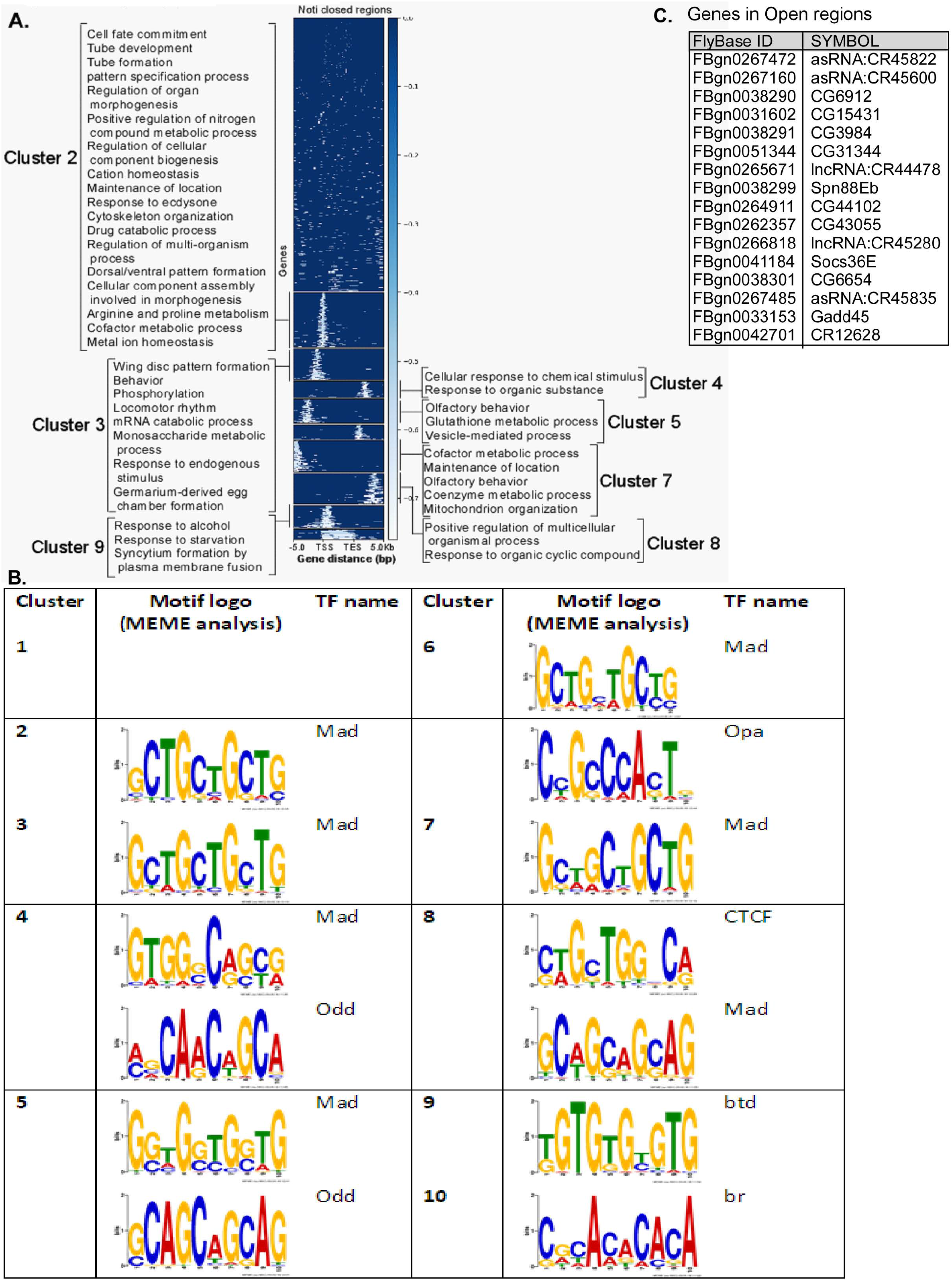
Analysis of changes in chromatin actability upon loss of Not. **(A)** Whole genome changes in chromatin accessibility unveiled by ATAC seq divided to clusters by location along gene regions and GO ontology of each cluster. **(B)** MEME analysis of cluster-enrichment in DNA binding sequences associated with the indicated TFs. **(C)** List of all genes in the vicinity of regions that exhibit increased accessibility upon loss of Non -stop in ECs.

**Figure 6; Supplemental Figure 1:**
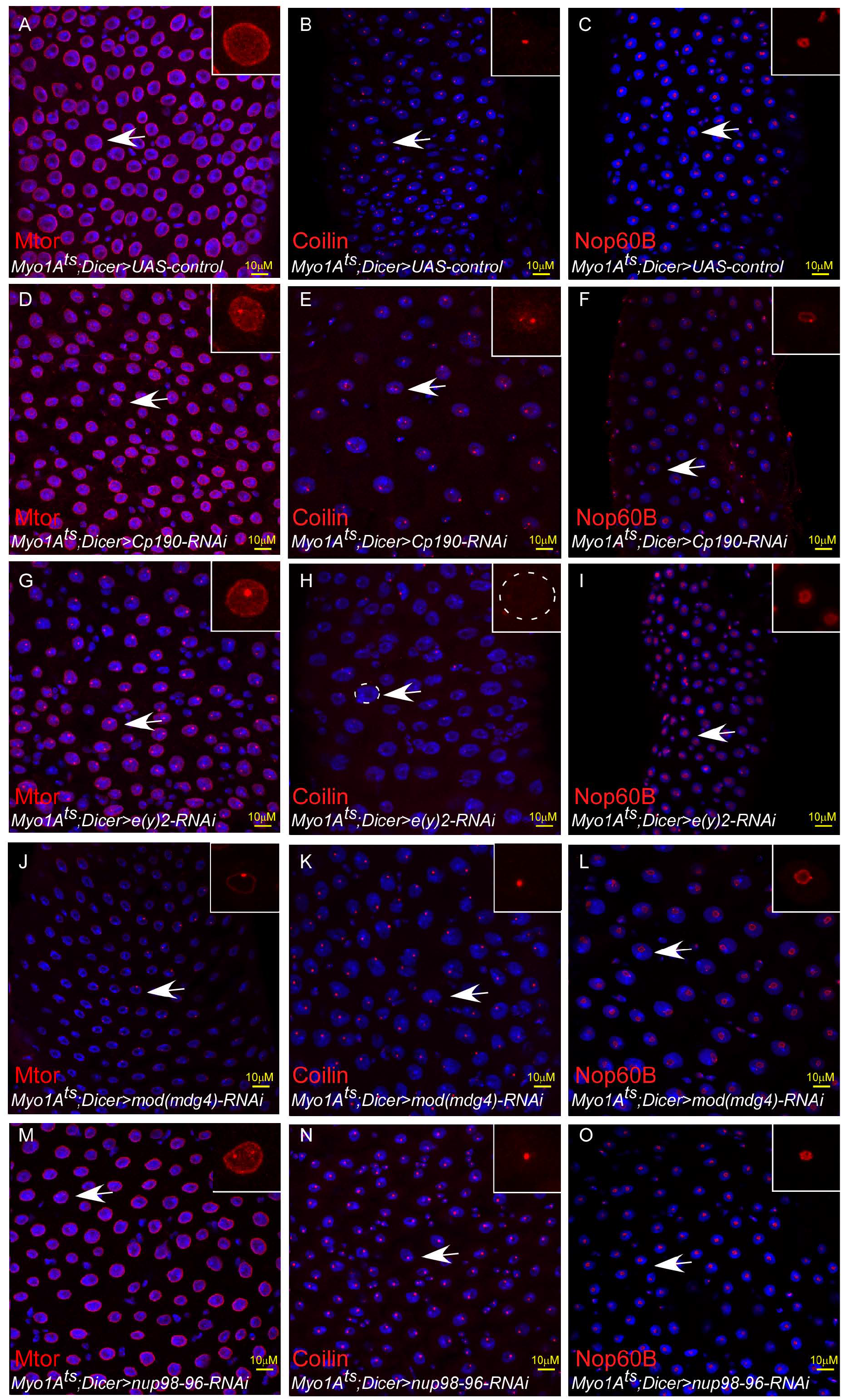
NIC subunits are required for maintaining large-scale organization of the EC nucleus. **(A-O)** Confocal images of the midgut tissue using the indicated antibodies (red) and expressing the indicated transgenes in EC using the MyoIA-Gal4/Gal80ts system, DAPI marks DNA (blue) and scale bar is 10μM M (A, D, G, J, M) Mtor; (B, E, H, K, N) Coilin (C, F, I, L, O) Nop60B ; (A-C) Control, (D-F) UAS-Cp190 RNAi; (G-I) UAS-e(y)2 RNAi (J-L) Mod(Mdg4) RNAi (M-O) Nup98-96 RNAi. **Figure 6; Supplemental source data**: Quantification of cell populations described in 6I-L.

**Figure 8 Supplemental Figure 1:**
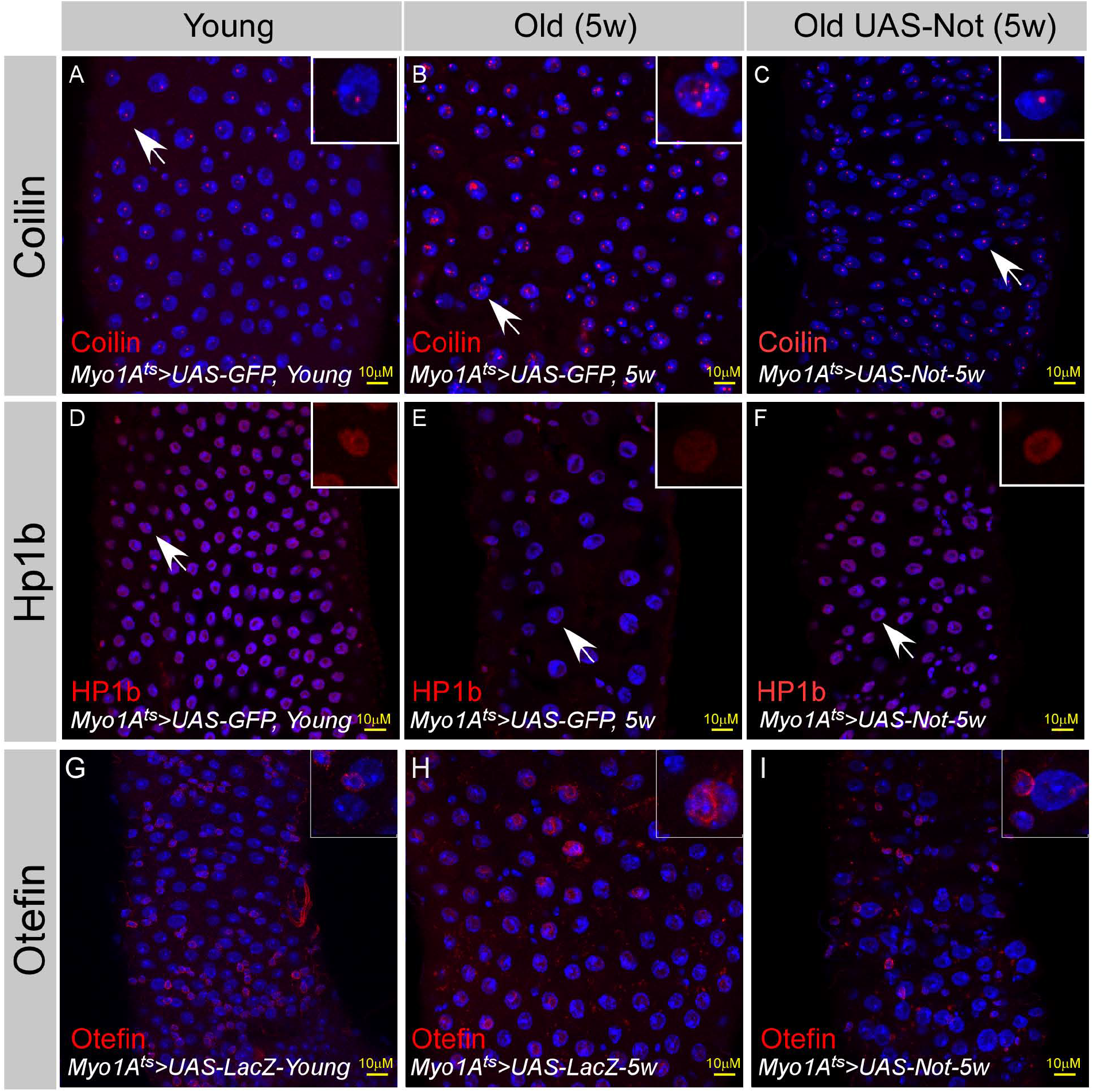
Expression of Non-stop restore large-scale organization of aged ECs: **(A-H)** Confocal images of the midgut tissue using the indicated antibodies and expressing the indicated transgenes in EC using the MyoIA-Gal4/Gal80ts (A, D, G) Young Guts expressing the indicated UAS-LacZ. Five weeks old guts expressing the indicated control. (C, F, H) Five weeks old guts expressing UAS-Non-stop. (A-C) a-Coilin; (D-F) a-HP1b; (G-I) a-Otefin.

## Supplemental Tables

### Key Resources table

**Table S1:** Results of primary and secondary transgenic RNAi screens

**Table S2:** Proteomic analysis of Non-stop bound proteins

**Table S3:** RNA-seq of Not-regulated genes

**Table S4:** ATAC-seq profiling of non-stop dependent changes in chromatin accessibility

**Table S5:** Gene clustering of Non-stop closed regions (complement Fig. 5 Figure Supp. 3)

## Materials and Methods

- Key resource table with fly stocks and antibodies used in this study
- Plasmids and Primers used in this study
- Chemicals used

### Methods

- In vitro binding
- Direct Yeast 2 Hybrid
- Proteomic analysis of Non-stop associated proteins
- RNAi in *Drosophila* S2 cells
- Conditional expression of transgenes in specific gut cells
- Conditional G-TRACE analysis
- Gut dissection and immunofluorescence detection
- Gut integrity and tracing of organismal survival
- Genomic analysis; RNA-seq, ATAC-seq and bioinformatics analyses including RNA extraction, cDNA preparation and Gene expression and RNA-seq, and bioinformatics analyses.
- Statistical analysis

### Fly stocks used in this study

Fly stocks were maintained on yeast-cornmeal-molasses-malt extract medium at 18°C or as stated in the text. UAS-RNAi used in the screen are described under Table S1.

### UAS and Gal4 transgenic lines used

All transgenic RNAi lines used for the Ub/Ubl screen are detailed in Table S1. All other lines used in this study are described in the

### Antibodies used in this study

All primary and secondary antibodies used are described in the

Key resource table.

### Plasmids and primers

pRmha3 C-HAx2-FLx2-nonstop-735 – was as described in (Cloud et al. 2019).

#### Plasmids for in vitro binding

CP190 CT (aa, 468-1096) was PCR-amplified using primers 5’-tttggtaccgggccctggctgtgcctg-3’ and 5’-tttctcgagtgcggccgcagatcttag-3’ and subcloned into pET32a(+) vector (Merck Biosciences) in frame with 6xHis tag using restriction sites *KpnI* and *XhoI.*

CP190 NT (aa, 1-524) was PCR-amplified using primers 5’ - tttcatatgggtgaagtcaagtccgtg −3’ and 5’-tttctcgagcatgtggaaatgcagttcccg −3’ and subcloned into pET32a(+) vector (Merck Biosciences) in frame with 6xHis tag using restriction sites *NdeI* and *XhoI.*

E(y)2 was PCR-amplified using primers 5’ - tttggatccccggaattcccgacgatgag-3’ and 5’-tttgcggccgcttaggattcgtcctctggc-3’ and subcloned into pET32a(+) vector (Merck Biosciences) in frame with 6xHis tag using restriction sites *BamHI* and *NotI*

#### Plasmids used in the yeast two-hybrid assay

The full-sized Not (aa 496) was PCR-amplified using primers 5’-ttgaattcatgtccgagacgggttgtc-3’ and 5’-ttgtcgacttactcgtattccagcacatt-3’ and subcloned into pGBT9 vector (Clontech) in frame with DNA-binding domain of GAL4 using restriction sites *EcoRI* and *Sal1*.

The full-sized CP190 (aa 1096) was PCR-amplified using primers 5’-ttcccgggcatgggtgaagtcaagtccg-3’ and 5’-tttggaggagctatatttactaagatct-3’ and subcloned into pGAD424 vector (Clontech) in frame with activation domain of GAL4 using restriction sites *SmaI* and *BamHI.* Fragments of CP190 from first to fourth zinc fingers was PCR-amplified using primers 5’-ttgaattcgagaatactactgggccct-3’ and 5’-ttgtcgacgccatcctccaaagcctg-3’, from second to third - 5’-ttgaattcgcgctttgtgagcattgc-3’ and 5’-ttgtcgacgttgtcgtccgtgtgcac-3’ and then subcloned into pGAD424 vector (Clontech) in frame with activation domain of GAL4 using restriction sites *EcoR1* and *Sall.*

Corresponding primers were used to make full-sized deletion variants of CP190:

CP 190Δ4 5’-aaggtaccggagcaggctttgga-3’ and 5’-aaggtacccactgctgcttgttgtcg-3’;

CP190Δ3-4 5’-aaggtaccggagcaggctttgga and 5’-aaggtaccaacgtatacagcagcgac-3’;

CP190Δ2-4 5’-aaggtaccggagcaggctttgga and 5’-aaggtacccgcgccggatcaattg-3’;

CP190Δ1-4 5’-gccctggctgaaggagcaggctttggagga and 5’-cctgctccttcagccagggcccagtagtat-3’,

#### Primers used for Non-stop RNAi in Drosophila cells

- Not-RNAi forward – 5’ -cggaattccgaattaatacgactcactatagggatttaatctggaaccatgcgaa-3’
- Not-RNAi reverse – 5’ -cggaattccgaattaatacgactcactatagggaaatgtcccaaaacggatcgta-3’

### Chemicals

Bromophenol Blue (Sigma #B5525), Guanidine hydrochloride (Sigma #G4505), NP40 (Igepal CA-630) (Sigma #I3021), Triton X-100 (Amresco #0694), Acrylamide (Bis-Acrylamide 29:1) (Biological Industries #01-874-1A), Ammonium Persulfate (Sigma #A-9164), TEMED (Sigma #T-7024), L-Glutamine (Gibco #25030024), MG132 (Boston Biochemicals), Blot Qualified BSA (Biological Industries #PRW3841), Agarose (SeaKem® LE Agarose-Cambrex Bio Science #CAM-50004), Bradford Protein Assay (BioRad #500- 0006), EZ-ECL (Biological Industries #20-500-500), FD&C blue dye #1, Cyclohexamide (Sigma #01810).

## Methods

### In vitro binding

6xHis-tagged proteins were xpressed and purification from E. coli BL-21 (DE3), using Ni-NTA agarose beads. His-tagged protein were induced with 0.5mM IPTG for 5h at 30°C and subsequently immobilized on with Ni-NTA agarose beads. Nuclear extract derived from Non-stop expressing S2 cells was prepared similar to the described in (Cloud et al, 2019), see “Non-denaturing extract”: Stably transfected cells were resuspended in Extraction Buffer (20 mM HEPES (pH7.5), 25% Glycerol, 420 mM NaCl, 1.5 mM MgCl_2_, 0.2 mM EDTA, 1:100 ethidium bromide with protease inhibitors added. 1% NP-40 was added and the cells were pipetted up and down until the solution was homogenous. They were placed on ice for one hour with agitation every 10-15 minutes. They were then centrifuged for 30 minutes at 4 °C at 20,000 x g. An equal volume of Dignum A buffer (10 mM HEPES (pH 7.5), 1.5 mM MgCl2, 10 mM KCl) was added to the lysates in order to adjust the salt concentration to 210 mM NaCl.

Binding was performed using 0.5mg of S2 cell extract expressing HF-Non-stop and the indicated His-tagged proteins immobilized to Ni-NTA beads using binding buffer (20mM Hepes-KOH pH 7.7, 150mM NaCl, 10mM MgCh0.1%mM ZnCl2, 0.1% NP40, 10% Glycerol and protease inhibitors) for over-night in rotation. Subsequently, beads were collected and washed four times with wash buffer, and proteins resolved over SDS-PAGE and detected by western blot analysis.

### Yeast two-hybrid assay (Y2H)

Y2H was carried out using yeast strain pJ69-4A (MATa trp1-901 leu2-3,112 ura3-52 his3-200 gal4Δ gal80Δ GAL2-ADE2 LYS2::GAL1-HIS3 met2::GAL7-lacZ), with plasmids according to Clontech protocols. In brief, for growth assays, AD (activation domain of GAL4) - and BD (DNA-binding domain of GAL4) -fused plasmids were co-transformed into yeast strain pJ69-4A by the lithium acetate method, as described by the manufacturer with some modifications. Transformed cells were plated on selective medium lacking Leu (leucine biosynthesis gene Leu2 is expressed from pGAD424 plasmid) and Trp (tryptophan biosynthesis gene Trp1 is expressed from pGBT9 plasmid) (‘medium-2’). The plates were incubated at 30°C for 2-3 days. Afterward, the colonies were streaked out on plates on selective medium lacking either Leu, Trp and His (histidine biosynthesis gene His3 is used as reporter in Y2H assay) (‘medium-3’). The plates were incubated at 30°C for 3-4 days, and growth was assessed. The positive growth of yeast on selective ‘medium-3’ indicates a physical interaction between protein molecules fused with AD and BD. Each assay was prepared as three independent biological replicates with three technical repeats.

### Proteomic analysis

Multidimensional protein identification technology and Mass spectrometry data processing were identical to the described in detail at (Cloud et al. 2019); Multidimensional protein identification technology (MudPIT) and Mass spectrometry data processing were identical to that described in Cloud et al. 2019. MudPIT: TCA-precipitated protein pellets were solubilized using Tris-HCl pH 8.5 and 8 M urea, followed by addition of TCEP (Tris(2-carboxyethyl)phosphine hydrochloride; Pierce) and CAM (chloroacetamide; Sigma) were added to a final concentration of 5 mM and 10 mM, respectively. Proteins were digested using Endoproteinase Lys-C at 1:100 w/w (Roche) at 37°C overnight. Samples were brought to a final concentration of 2 M urea and 2 mM CaCl2 and a second digestion was performed overnight at 37°C using trypsin (Roche) at 1:100 w/w. The reactions were stopped using formic acid (5% final). The digested size exclusion eluates were loaded on a split-triple-phase fused-silica micro-capillary column and placed in-line with a linear ion trap mass spectrometer (LTQ, Thermo Scientific), coupled with a Quaternary Agilent 1100 Series HPLC system. The digested Non-stop and control FLAG-IP eluates were analyzed on an LTQ-Orbitrap (Thermo) coupled to an Eksigent NanoLC-2D. In both cases, a fully automated 10-step chromatography run was carried out. Each full MS scan (400-1600 m/z) was followed by five data-dependent MS/MS scans. The number of the micro scans was set to 1 both for MS and MS/MS. The settings were as follows: repeat count 2; repeat duration 30 s; exclusion list size 500 and exclusion duration 120 s, while the minimum signal threshold was set to 100. Mass Spectrometry Data Processing: The MS/MS data set was searched using ProLuCID (v. 1.3.3) against a database consisting of the long (703 amino acids) isoform of non-stop, 22,006 non-redundant Drosophila melanogaster proteins (merged and deduplicated entries from GenBank release 6, FlyBase release 6.2,2 and NCI RefSeq release 88), 225 usual contaminants, and, to estimate false discovery rates (FDRs), 22,007 randomized amino acid sequences derived from each NR protein entry. To account for alkylation by CAM, 57 Da were added statically to the cysteine residues. To account for the oxidation of methionine to methionine sulfoxide, 16 Da were added as a differential modification to the methionine residue. Peptide/spectrum matches were sorted and selected to an FDR less than 5% at the peptide and protein levels, using DTASelect in combination with swallow, an in-house software.

The permanent URL to the dataset is: ftp://massive.ucsd.edu/MSV000082625. The data is also accessible from: ProteomeXChange accession: PXD010462 http://proteomecentral.proteomexchange.org/cgi/GetDataset?ID=PXD010462. MassIVE | Accession ID: MSV000082625 - ProteomeXchange | Accession ID: PXD010462.

### RNAi in S2 cells

S2 Schneider DRSC cells (*Drosophila* Genomics Resource Center #181, RRID:<u>CVCL_Z992</u>) were maintained in Schneider’s media supplemented with 10% fetal bovine serum and 1% penicillin-streptomycin (Thermo-Fisher, Catalog number: 15070063, 5000 U/ml) RNAi in S2 cells was performed as described in (Abed et al. 2011).

### Conditional expression of transgenes in specific gut cells

Conditional expression of transgenic lines in specific midgut cells was achieved by activating a UAS-transgene under the expression of the cell-specific Gal4-drivers together with the tubGal80^ts^ construct (Jiang et al., 2009). Flies were raised at 18°C. 2-4 days old, F1 adult progeny were transferred to the restrictive temperature 29°C (Gal80 off, Gal4 on) for two days unless indicated otherwise, dissected and analyzed. At least three biological independent repeats were performed for each experiment. Where possible, multiple RNAi lines were used.

### Conditional G-TRACE analysis

G-TRACE analyses was as described in (Flint-Brodsly et al. 2019) using Myo-Gal4; G-TRACE flies were crossed to UAS-LacZ; Gal80^ts^ (control) or UAS-Non-stop RNAi ; Gal80^ts^ and the appropriate genotypes were raised at 18°C (a temperature where no G-TRACE signal was detected). At 2-4-days, adult females were transferred to 29°C and linage tracing was performed.

### Gut dissection and immunofluorescence detection

Gut fixation and staining were carried out as previously described (Shaw et al., 2010; Flint-Brodsly et al. 2019).

### Gut integrity and animal survival

Young female flies from the indicated genotype were collected into a fresh vial (10 flies per vial), that were kept in a humidified, temperature-controlled incubator at 29°C for the indicated time period. Smurf assay was performed as described in (Flint-Brodsly et al. 2019). Flies were transferred into vials containing fresh food every two days and were scored for viability at the indicated time points. LT50 (lethal time in days at which 50% of the flies died) Statistical analysis was calculated using the GraphPad Prism 5.00 (GraphPad Software, San Diego, CA, USA).

### Genomic studies

#### RNA-sequencing

RNA Sample Preparation: RNA-seq was performed similar to the described in (Flint-Brodsly et al 2019). In brief, Adult *Drosophila* (2-4 days old) females, from 4 biological repeats, in which UAS-Non-stop RNAi or control UAS-GFP RNAi were expressed in ECs using MyoIAts and dissected in Ringer’s solution on ice. The solution was then discarded, and the guts were disrupted by adding 350μl RLT+β ME buffer (350μl RLT+ 3.5μl β-ME). Guts were than vortex for homogenization. 350 μl of 70% ethanol was then added and mixed well by pipetting. Guts were uploaded into RNeasy spin column and RNA purified according to the manufacture instructions. Sample quality (QC) Quality measurements for total RNA were performed using the TapeStation 2200 (Agilent).

#### Library preparation and data generation of RNA-sequencing

Eight RNA-seq libraries were produced using the NEBNext® Ultra Directional RNA Library Prep Kit for Illumina (NEB, cat no. E7420) according to manufacture protocol and starting with 100 ng of total RNA. mRNA pull-up was performed using the Magnetic Isolation Module (NEB, cat no. E7490). Two out of the twelve libraries (samples B1 & B2) were disqualified based on low library yield and high levels of adaptor dimer. The remaining ten libraries were mixed into a single tube at an equal molar concentration. The RNA-seq data was generated on two lanes of HiSeq2500, 50SR.

NGS QC, alignment and counting 50 bp single-end reads were aligned to *Drosophila* reference genome and annotation file (*Drosophila* melanogaster BDGP6 downloaded from ENSEMBL) using TopHat (v2.0.13) allowing 2 mismatches per read with options -very-sensitive. The number of reads per gene was counted using Htseq (0.6.0).

#### Descriptive and RNA-seq DEGs analysis

Samples’ clustering and differential expressed genes (DEGs) were calculated using Deseq2 package (version 1.10.1). The similarity between samples was evaluated using correlation matrix, shown a heat plot and Principal Component Analysis (PCA). Samples belonging to the same group were more similar then samples from different experimental groups (Figure 5 Figure Supplemental 1A). The expression ~12,000 fly transcripts were compared using DESeq2 and list of the differentially express genes (DEGs) was extracted into excel files. At adjusted p-value (p-adj) &lt;0.01 and LogFC &gt; 1 or LogFC &lt; −1, 1428 DEGs were found between guts derived from the control vs Non-stop RNAi targeted EC. Moreover, the expression of Not was found to be downregulated by −0.88 log2FC between Non-stop RNAi vs. Ctrl with p-adj of less than 10-9.

#### ATAC (Assay for Transposase-Accessible Chromatin) sequencing

Adult *Drosophila* (2-4 days old) females, from 3 biological repeats, in which Non-stop RNAi or GFP RNAi control were expressed in ECs using MyoIAts were dissected in ice cold Ringer’s solution, and immediately placed in 25μl of ice cold ATAC lysis buffer (10 mM Tris-HCl, pH 7.4,10mM NaCl, 3mM MgCl2, 0.1% IGEPAL CA-630). Lysed guts were then centrifuged at 500xg for 15 minutes at 4’C and the supernatant was discarded. The rest of the ATAC-seq protocol was performed as described in (Buenrostro et al., 2013). The final library was purified using a Qiagen MinElute kit (Qiagen) and Ampure XP beads (Ampure) (1:1.2 ratio) were used to remove remaining adapters. All samples were quantified using Qubit DNA HS assay. The final library was first checked on an Agilent Bioanalyzer 2000 for quality and the average fragment size. Successful libraries were sequenced with NextSeq 75 cycles high-output flow-cell, targeting ~25 million reads/sample.

#### Bioinformatic analysis of ATAC-seq

Raw reads were trimmed for adapters and aligned to the Drosophila melanogaster reference genome using bowtie2. Redundant duplicated reads that aligned to the exact locations were removed from the aligned results, and then converted to the tagAlign format with consideration of the strand shift (“+” strand reads shifted by 4bp, and strand reads shifted by −5bp). The tag-Align format alignment results were used to call peaks using MAC2. Narrow peaks with p<0.1 were reported. Peaks were compared across biological duplicates and pseudo duplicates (i.e., random subsets from a sample) to get IDR peaks that are supposed to be consensus across duplicates. Test for the difference between the two groups (control & Noti) has been performed with the Bioconductor R package DiffBind. Significance is set by FDR<0.1.

#### Comparisons between RNA and ATAC sequencing data

The two data sets of DEGs and DBAs were analyzed to create a list of significant differentially bound peaks that are close to genes from the top 1428 DE genes. The thresholds that have been used to associate the peaks to genes is within 10kb upstream and 10kb downstream of the genes. To look for common binding motives between the two data sets, the genes were sub-divided into 4 categories: 1. peaks overlapping promoters. 2. peaks upstream of promoters. 3. peaks in genic region. 4. peaks downstream of genes. Next, I performed motif finding informatics using CONSENSUS, MDScan and MEME software.

Gene ontology analysis of mRNA expression and ATAC-seq was performed using the Database for Annotation, Visualization and Integrated Discovery (DAVID http://david.abcc.ncifcrf.gov/home.jsp; v6.7 and v6.8) using default settings (2-fold p<0.05 Benjamini P value for analysis E-03) (Huang et al., 2009).

Biological processes with a p-value lower than 0.05 were further analyzed with Revigo (Supek F et al., 2011). Gene ontology analyses via Cytoscape: ClueGO app (v2.2.5) in Cytoscape (v 3.4.0) was used to conduct GO enrichment analyses. In our study, ClueGO was used to identify different functional groups in the following terms: Biological Process (BP), Cellular Component (CC) and Molecular Function (MF) enrichment analysis. A p-value ≤ 0.05 was used as the cut-off criterion.

### Statistical analysis

Data was collected from three independent experiments. Statistical analysis, z-test comparisons were performed using Prism6 ANOVAs software. Significance is indicated by *** = P<0.001 and ** = P<0.01.

